# Transcription factor RFX7 governs a tumor suppressor network in response to p53 and stress

**DOI:** 10.1101/2021.03.25.436917

**Authors:** Luis Coronel, Konstantin Riege, Katjana Schwab, Silke Förste, David Häckes, Lena Semerau, Stephan H. Bernhart, Reiner Siebert, Steve Hoffmann, Martin Fischer

## Abstract

Despite its prominence, the mechanisms through which the tumor suppressor p53 regulates most genes remain unclear. Recently, the regulatory factor X 7 (RFX7) emerged as a suppressor of lymphoid neoplasms, but its regulation and target genes mediating tumor suppression remain unknown. Here, we identify a novel p53-RFX7 signaling axis. Integrative analysis of the RFX7 DNA binding landscape and the RFX7-regulated transcriptome in three distinct cell systems reveals that RFX7 directly controls multiple established tumor suppressors, including PDCD4, PIK3IP1, MXD4, and PNRC1, across cell types and is the missing link for their activation in response to p53 and stress. RFX7 target gene expression correlates with cell differentiation and better prognosis in numerous cancer types. Interestingly, we find that RFX7 sensitizes cells to Doxorubicin by promoting apoptosis. Together, our work establishes RFX7’s role as a ubiquitous regulator of cell growth and fate determination and a key node in the p53 transcriptional program.

## Introduction

RFX7 belongs to a family of eight transcription factors that share a highly conserved DNA-binding domain (DBD) through which they can bind to *cis*-regulatory X-box motifs (1, 2). *RFX5* is the closest sibling of *RFX7*, and while the expression of most *RFX* genes is restricted to specific cell types, *RFX1*, *RFX5*, and *RFX7* display ubiquitous expression (2, 3). Whole-genome sequencing efforts led us and others to discover *RFX7* mutations in 13 to 15 % of Epstein-Barr Virus-negative Burkitt lymphoma patients (4, 5). Additionally, genome-wide association studies linked *RFX7* to chronic lymphocytic leukemia (6–8). *RFX7* alterations have also been identified in diffuse large B cell lymphoma (9), acute myeloid leukemia (10), as well as in mouse models of lymphoma (9, 11) and leukemia (12). In addition to hematopoietic neoplasms, *RFX7* has been associated with body fat distribution (13), Alzheimer’s disease (14), and autism spectrum disorder (15), suggesting that RFX7 may function in various cell types and tissues. While human RFX7 is functionally uncharacterized, first insights from animal models identified Rfx7 to play a role in anuran neural development (16) and maturation and metabolism in murine lymphoid cells (17). Importantly, the regulation of RFX7 and its target genes mediating tumor suppression are unknown.

In response to stress conditions, p53 transcriptionally regulates a plethora of target genes to suppress tumorigenesis (18, 19). Thereby, p53 influences diverse cellular processes, including apoptosis, cell cycle progression, and metabolism. Using integrative omics approaches, we started to disentangle the p53 gene regulatory network (GRN) into subnetworks of genes controlled directly by p53 or indirectly through downstream transcription factors (20). For example, p53 regulates the largest subset of genes indirectly through its direct target gene *CDKN1A*, encoding cyclin-dependent kinase inhibitor p21 and reactivating DREAM and RB:E2F *trans*-repressor complexes to down-regulate cell cycle genes (20–23). However, complex cross-talks between signaling pathways impede the identification of indirect regulations. Uncovering the molecular mechanisms through which p53 indirectly controls most p53-regulated genes, therefore, remains a longstanding challenge (19).

Our findings place the understudied transcription factor RFX7 immediately downstream of p53 and provide compelling evidence for RFX7’s ubiquitous role in governing growth regulatory pathways. We reveal that RFX7 orchestrates multiple established tumor suppressor genes in response to cellular stress. Thus, RFX7 emerges as a crucial regulatory arm of the p53 tumor suppressor. In the context of cancer biology, the general importance of this new signaling axis is exemplified by the better prognosis of patients with a medium to high expression of RFX7 targets across the TCGA pan-cancer cohort, which indicates recurrent de-regulation of RFX7 signaling in cancer.

## Materials & Methods

### Cell culture, drug treatment, and transfection

U2OS and HCT116 cells (ATCC, Manassas, Virginia, USA) were grown in high glucose Dulbecco’s modified Eagle’s media (DMEM) with pyruvate (Thermo Fisher Scientific, Darmstadt, Germany). RPE-1 hTERT cells (ATCC) were cultured in DMEM:F12 media (Thermo Fisher Scientific). Culture media were supplemented with 10% fetal bovine serum (FBS; Thermo Fisher Scientific) and penicillin/streptomycin (Thermo Fisher Scientific). Cell lines were tested twice a year for *Mycoplasma* contamination using the LookOut Detection Kit (Sigma), and all tests were negative.

Cells were treated with DMSO (0.15 %; Carl Roth, Karlsruhe, Germany), Nutlin-3a (10 µM; Sigma Aldrich, Darmstadt, Germany), Actinomycin D (5 nM; Cayman Chemicals, Ann Arbor, Michigan, USA), or Doxorubicin (1 µM or as indicated; Cayman Chemicals) for 24 h. For knockdown experiments, cells were seeded in 6-well plates or 6 cm dishes and reverse transfected with 5 nM Silencer Select siRNAs (Thermo Fisher Scientific) using RNAiMAX (Thermo Fisher Scientific) and Opti-MEM (Thermo Fisher Scientific) following the manufacturer protocol.

Images of cells were taken using an Evos M5000 microscope (Thermo Fisher Scientific) or a ChemiDoc MP documentation system (Bio-Rad, Feldkirchen, Germany).

### Chromatin immunoprecipitation, RNA extraction, and reverse transcription semi-quantitative real-time PCR (RT-qPCR)

ChIP was performed with the SimpleChIP Kit (Cell Signaling Technology, Canvers, MA, USA) following the manufacturer instructions. 3 µg of p53 (kind gift from Dr. Bernhard Schlott (24)) or RFX7 (#A303-062A Bethyl Laboratories, Montgomery, TX, USA) antibody were used per IP. Sonication was performed on a Bioruptor Plus (Diagenode, Seraing, Belgium).

Total cellular RNA was extracted using the RNeasy Plus Mini Kit (Qiagen, Hilden, Germany) following the manufacturer protocol. One-step reverse transcription and real-time PCR was performed with a Quantstudio 5 (Thermo Fisher Scientific) using Power SYBR Green RNA-to-CT 1-Step Kit (Thermo Fisher Scientific) following the manufacturer protocol. We identified *ACTR10* as a suitable control gene that is not regulated by p53 but expressed across 20 gene expression profiling datasets (20). Generally, two or three biological replicates with three technical replicates each were used. Given the nature of the technical setup, a few individual data points were erroneous and, thus, excluded (see source data).

Primer sequences are listed in Supplementary Table 4.

### Western blot analysis

Cells were lysed in RIPA buffer (Thermo Fisher Scientific) containing protease and phosphatase inhibitor cocktail (Roche, Grenzach-Wyhlen, Germany or Thermo Fisher Scientific). Protein lysates were scraped against Eppendorf rack for 20 times and centrifuged with 15000 rpm for 15 min at 4°C. The protein concentration of supernatant lysates was determined using the Pierce 660 nm Protein Assay Kit (Thermo Fisher Scientific) and a NanoDrop1000 Spectrophotometer (Thermo Fisher Scientific). Proteins were separated in a Mini-Protean TGX Stain-Free Precast 4-15% Gel (Bio-Rad) using Tris/Glycine/SDS running buffer (Bio-Rad). Proteins were transferred to a 0.2 µm polyvinylidene difluoride (PVDF) transfer membrane either using a Trans-Blot Turbo Mini Transfer Pack (Bio-Rad) in a Trans-Blot Turbo (Bio-Rad) or using a Mini Trans-Blot Cell (Bio-Rad) in a Mini-Protean Tetra Cell (Bio-Rad). Following antibody incubation, membranes were developed using Clarity Max ECL (Bio-Rad) and a ChemiDoc MP imaging system (Bio-Rad).

Antibodies and their working concentrations are listed in Supplementary Table 4.

### Pre-processing of Illumina sequencing data

Quantification and quality check of libraries were performed using the Agilent Bioanalyzer 2100 in combination with the DNA 7500 Kit. Libraries were pooled and sequenced on a NextSeq 500 (75 bp, single-end), HiSeq 2500 (50 bp, single-end), and NovaSeq 6000 (S1 or SP, 100 cycles). Sequence information was extracted in FastQ format using Illumina’s bcl2FastQ v2.19.1.403 or v2.20.0.422.

We utilized Trimmomatic (25) v0.39 (5nt sliding window approach, mean quality cutoff 22) for read quality trimming according to inspections made from FastQC (https://www.bioinformatics.babraham.ac.uk/projects/fastqc/) v0.11.9 reports. Illumina universal adapter as well as mono-and di-nucleotide content was clipped using Cutadapt v2.10 (26). Potential sequencing errors were detected and corrected using Rcorrector v1.0.3.1 (27). Ribosomal RNA (rRNA) transcripts were artificially depleted by read alignment against rRNA databases through SortMeRNA v2.1 (28). The preprocessed data was aligned to the reference genome hg38, retrieved along with its gene annotation from Ensembl v.92 (29), using the mapping software segemehl (30, 31) v0.3.4 with adjusted accuracy (95%) and split-read option enabled (RNA-seq) or disabled (ChIP-seq). Mappings were filtered by Samtools v1.10 (32) for uniqueness and properly aligned mate pairs. We removed duplicated reads with Picard MarkDuplicates v2.23.4.

### ChIP-seq and analysis

ChIP was performed as described above in biological duplicates for RFX7 ChIP and input DNA from Nutlin-3a and DMSO control treated U2OS, HCT116, and RPE-1 cells. Libraries were constructed using the NEBNext Ultra II DNA Library Preparation Kit (New England Biolabs, Frankfurt am Main, Germany) following the manufacturer’s description. Following pre-processing of the sequencing data (see above), biological replicates of each input and IP were pooled prior to peak calling with MACS2 v2.2.7.1 (33) with q-value cutoff 0.05. MACS2 was executed in both available modes utilizing either the learned or a customized shifting model parameterized according to the assumed mean fragment length of 150bp as extension size. The resulting peak sets were merged by overlap with BEDTools v2.29.2 (34). Per interval, the strongest enrichment signal under the associated peak summits as well as the lowest p-value and q-value was kept. ENCODE blacklist regions (35) were filtered out. Unique and shared overlapping peak sets were identified using BEDTools ‘intersect’. *De novo* motif discovery was performed using ‘findMotifsGenome’ of HOMER v4.10 (36) with options *-size given -S 15*. The top X-box motif recovered from the *de novo* analysis of the 120 overlap peaks with relaxed log odds detection threshold of 7 was used to discover X-boxes across hg38 using HOMER’s ‘scanMotifGenomeWide’. Conservation plots displaying the average vertebrate PhastCons score (37) were generated using the Conservation Plot tool in Cistrome (38). The Cis-regulatory Element Annotation System (CEAS) tool in Cistrome (38) was used to identify the enrichment of binding sites at genome features. Genes associated with RFX7 peaks were identified using BETA-minus in Cistrome (38) with a threshold of 5 kb from the TSS. To identify whether RFX7 functions as an activator or repressor of gene transcription, we employed BETA analysis (39) in Cistrome (38). CistromeDB toolkit (40) was used to identify TFs that display ChIP-seq peak sets (top 10k peaks) that are significantly similar to the set of 120 common RFX7 peaks. Bigwig tracks were generated using deeptools ‘bamCoverage’ with options –binSize 1 and –extendReads 150 (41).

Publicly available p53 ChIP-seq data from Nutlin-3a-treated U2OS (42) and HCT116 (43) cells was obtained from CistromeDB (40). Ten publicly available RFX5 ChIP-seq datasets from A549, GM12878, HepG2, hESC, IMR90, K562, MCF-7, HeLa, and SK-N-SH cells were obtained from CistromeDB and joined using BEDTools ‘multiinter’ followed by ‘merge’. RFX5 peaks supported by at least 5 out of the 10 datasets were kept for further analyses.

### RNA-seq and analysis

Cellular RNA was obtained as described above in biological triplicates or quadruplets. Quality check and quantification of total RNA were performed using the Agilent Bioanalyzer 2100 in combination with the RNA 6000 Nano Kit (Agilent Technologies). Libraries were constructed from 1 µg of total RNA using Illumina’s TruSeq stranded mRNA Library Preparation Kit or from 500 ng total RNA using NEBNext Ultra II RNA -polyA+ (mRNA) Library Preparation Kit (New England Biolabs) following the manufacturer’s description.

Following pre-processing of the data (see above), read quantification was performed on exon level using featureCounts v1.6.5 (44), parametrized according to the strand specificity inferred through RSeQC v3.0.0 (45). Differential gene expression and its statistical significance was identified using DESeq2 v1.20.0 (46). Given that all RNA-seq data was derived from PolyA-enriched samples, we only included Ensembl transcript types ‘protein_coding’, ‘antisense’, ‘lncRNA’, and ‘TEC’ in our analysis. Common thresholds for |log_2_(fold-change)| ≥ 0.25 and adj. p-value < 0.01 were applied to detect significant differential expression. Publicly available RNA-seq data from human p53-negative HL-60 promyelocytes differentiating into macrophages or neutrophils was obtained from GEO accession number GSE79044 (47). Publicly available RNA-seq data from of human umbilical cord blood-derived unrestricted somatic stem cells (USSC) differentiating into neuronal-like cells was obtained from GEO accession number GSE96642 (48). Publicly available RNA-seq data from human pluripotent stem cells differentiating into lung alveolar cells was obtained from GEO accession number GSE96642 (49). RNA-seq data from human cells were processed as described above. Publicly available RNA-seq data from conditional Rfx7 knock-out mice was obtained from GEO accession number GSE113267 (17). The mouse RNA-seq data was processed as described above, but aligned to the mouse reference genome mm10. Given the naturally larger variation in tissue samples, thresholds for |log_2_(fold-change)| ≥ 0.25 and adj. p-value ≤ 0.05 were applied to detect significant differential expression.

### p53 Expression Score

The *p53 Expression Score* has been published in a previous meta-analysis (20) and reflects a summary of p53-dependent gene expression from 20 genome-wide p53-dependent gene expression profiling datasets. In each dataset a gene was identified either as significantly down-regulated (score -1), significantly up-regulated (score +1), or not significantly regulated (score 0) by p53. The *p53 Expression Score* displays for each gene the sum of the scores from all 20 datasets in the meta-analysis.

### Transcription factor binding and motif enrichment analysis

We used iRegulon (50) to identify transcription factors and motifs that are enriched within 500 bp upstream of the TSS or within 10 kb around the TSS of selected genes.

### Cell viability data from the Cancer Dependency Map (DepMap) project

The DepMap project pursued a systematic knockdown of genes in a large panel of cancer cell lines to identify genes that are essential for cancer cell viability (51). RFX7 data was available for 343 cell lines in which RFX7 was depleted by RNAi (depmap.org). The DEMETER2 score is a dependency score that reflects the effect of a given knockdown on cell viability (52). Negative dependency scores reflect decreased cell viability upon loss of the target gene, while positive scores indicate increased cell viability.

### Cell proliferation and viability assay

U2OS and HCT116 were transfected with 5 nM of respective siRNAs using RNAiMAX. The next day, cells were seeded in 96-well plates (9 000 cells per well). After 24h of transfection, cells were treated with Doxorubicin or DMSO control for 24h. Subsequently, the cells recovered for 6 days in fresh drug-free media. WST-1 reagent (Sigma Aldrich) was added for 4 h following the manufacturer protocol before absorbance was measured at 440 nm on a M1000pro microplate reader (Tecan, Männedorf, Switzerland).

### Clonogenic assay

HCT116 cells were transfected with 5 nM of respective siRNAs using RNAiMAX. The next day, the transfected cells were seeded 6-well plates (50 000 cells per well) containing 2 ml of culture media. After 24h transfection, cells were challenged with ether DMSO or treated with different Doxorubicin concentrations (0.05 µM, 0.075 µM, 0.1 µM, 0.15 µM and 0.2 µM for 24 hrs. All plates were then recovered in drug-free media and growth continued for another 7 days. After 7 days of recovery, cells were stained with crystal violet containing glutaraldehyde solution and briefly rinsed with plain water.

### Annexin V assay

HCT116 cells were transfected with 5 nM of respective siRNAs using RNAiMAX. The next day, the transfected cells were seeded 6-well plates (50 000 cells per well) containing 2 ml of culture media. After 24h transfection, cells were challenged with ether DMSO or treated with different Doxorubicin concentrations (0.05 µM, 0.075 µM, 0.1 µM, 0.15 µM and 0.2 µM for 24 hrs. All plates were then recovered in drug-free media and growth continued for another 6 days. Cells were stained with Annexin V and PI using the Annexin V Apoptosis Detection Kit I (BD Biosciences, San Jose, CA, USA) following the manufacturer instructions. Cell staining was quantified through flow cytometry on a BD FACSAria Fusion (BD Biosciences) and flow cytometry data was analyzed using FACSDiva 9.0.1 (BD Biosciences).

### Survival analysis

Survival analyses for Cancer Genome Atlas (TCGA) cases were based on the expression of a set of 19 direct RFX7 targets. Specifically, genes in this set were required to have been identified in all three cell line models (Fig. 2a, Extended Data Fig. 2a) and to have a *p53 Expression Score* > 5 to avoid the inclusion of cell cycle genes and to filter for a reproducibly strong p53-RFX7 signaling response. This 19-gene-set comprises *TP53INP1*, *PNRC1*, *MXD4*, *PIK3IP1*, *TOB1*, *PIK3R3*, *SESN3*, *YPEL2*, *PLCXD2*, *SLC43A2*, *CCND1*, *IP6K2*, *TSPYL2*, *RFX5*, *PDCD4*, *CCNG2*, *ABAT*, *TSPYL1*, and *JUNB*. We retrieved clinical data and FPKM normalized gene expression values from TCGA using the R package TCGAbiolinks v2.18.0 (53). For the whole pan-cancer set and for each of the 33 cancer types we calculated single-sample expression scores for the 19-gene-set from FPKM transformed quantification data using the official GenePattern codebase v10.0.3 for single sample gene set enrichment analysis (ssGSEA; https://github.com/GSEA-MSigDB/ssGSEA-gpmodule) (54). A single-sample expression score measures the degree of coordinated up or down-regulation of genes in the given set. Subsequently, we subdivided the expression scores into three equally sized categorial groups (high, medium, low). Kaplan-Meier plots and multivariate Cox regression analysis based on the expression groups were performed on clinical time to event and event occurrence information using the R survival package v3.2–3. The Cox proportional hazards (PH) model was used to investigate the relation of patient survival and categorical expression levels. To control for confounding factors, gender and age were included into all models. In case of the pan-cancer cohort, we further included cancer type into the regression analysis. The rates of occurrence of events over time were compared between the groups using the fitted PH model. Additionally, confounding factors, the distribution of gender, age, and cancer type were visualized for each categorial group.

### Statistics

ChIP and RT-qPCR data was analyzed using a two-sided unpaired t-test. Cell viability data from WST-1 assays were analyzed using a Sidak-corrected two-way ANOVA test. Mean Z-scores were compared using a two-sided paired t-test. Violin plots display the median. Bar graphs display mean and standard deviation. *, **, ***, and n.s. indicate p-values <0.05, <0.01, <0.001, and >0.05, respectively. The number of replicates is indicated in each Figure legend. FDR from RNA-seq data were obtained from DESeq2 analysis (‘padj’ values). p-values from ChIP-seq data were obtained from MACS2 analysis. The experiments were not randomized and investigators were not blinded to allocation during experiments.

### Data availability

Sequencing data is accessible through GEO (55) series accession numbers GSE162157, GSE162158, GSE162159, GSE162160, GSE162161, GSE162163. Previously published RNA-seq data was obtained from GEO accession numbers GSE113267 (17), GSE79044 (47), GSE144464 (48), and GSE96642 (49). Previously published p53 ChIP-seq data was obtained from CistromeDB (40) IDs 82544 (43) and 33077 (42). Similarly, previously published RFX5 ChIP-seq data (56) was obtained from CistromeDB IDs 45649, 45692, 45730, 45823, 45863, 45893, 46037, 100232, 100233, 100797. Source data for Figures are available from the corresponding authors upon request.

## Results

### The p53 target RFX7 mediates gene activation and markedly differs from RFX5

To identify novel nodes in the p53 GRN, we performed an enrichment analysis for transcription factor binding to genes frequently up-regulated by p53 activation but not directly bound by p53. We focused on proximal promoters, as these are more likely to confer robust gene regulation across cell types. An analysis of publicly available ChIP-seq datasets revealed multiple hits indicating enriched RFX5 binding to the genes’ proximal promoters (Figure 1A). Given that the RFX family shares a conserved DBD and ChIP-seq data is publicly available only for RFX1 and RFX5, we initially included all RFX family members in our investigation. To elucidate the potential role of RFX transcription factors in the p53 GRN, we analyzed published p53-dependent gene expression data (20). We identified *RFX5* and *RFX7*, but no other RFX family member, as being frequently up-regulated by p53 (Figure 1B). Investigation of published p53 DNA binding data revealed that *RFX7* contains two p53 binding sites in the first intron (intron1), while other family members, such as *RFX5* and *RFX1*, did not display p53 binding (Figure 1C and Supplementary Figure 1A). To test whether RFX7 or its ubiquitously expressed siblings RFX5 and RFX1 (3) affect p53-dependent up-regulation of genes, we selected potential target genes out of the 1081 genes potentially up-regulated indirectly by p53 that were frequently identified to bind RFX5 (Figure 1A). We selected *PDCD4*, *PIK3IP1*, *MXD4*, and *PNRC1* that are frequently up-regulated by p53 and that were identified in all six RFX5 ChIP-seq tracks (Figure 1A). Notably, *PDCD4*, *PIK3IP1*, *MXD4*, and *PNRC1* encode established tumor suppressors, which have not yet been established as p53-responsive genes (57–60). To this end, we employed the osteosarcoma cell line U2OS, which possesses intact p53 and is frequently used to study p53 and its signaling pathway (20). To specifically activate p53, we pharmacologically inhibited MDM2, the central gatekeeper of p53 activity, using the small molecule Nutlin-3a (61). RT-qPCR data confirmed that *PDCD4*, *PIK3IP1*, *MXD4*, and *PNRC1* are up-regulated in response to Nutlin-3a treatment. Importantly, the Nutlin-3a-induced up-regulation of *PDCD4*, *PIK3IP1*, *MXD4*, and *PNRC1* was attenuated upon knockdown of p53 and RFX7 (Figure 1D). In contrast to RFX7, depletion of RFX1 and RFX5 did not affect the p53-dependent induction of *PDCD4*, *PIK3IP1*, *MXD4*, and *PNRC1*. Thus, despite its similarity to RFX5, RFX7 plays a clearly distinct and strikingly consequential role in regulating those genes. Significantly, p53-dependent up-regulation of *CDKN1A* was not affected by RFX7 depletion, providing further evidence that RFX7 functions downstream of p53. Intriguingly, *RFX5* also appeared to be up-regulated by this novel p53-RFX7 signaling axis (Figure 1D). Western blot analyses indicated a p53-dependent induction of RFX7 protein levels upon Nutlin-3a treatment. In particular, a lower migrating form of RFX7 was induced in response to p53 activation (Figure 1E). Moreover, protein levels of PDCD4 and PIK3IP1 followed the p53-RFX7-dependent up-regulation of their mRNAs (Figure 1E). Further, ChIP-qPCR data revealed that RFX7 occupies the promoter regions of *PDCD4*, *PIK3IP1*, *MXD4*, and *PNRC1*. Upon Nutlin-3a treatment, RFX7 occupancy increased, while p53 did not occupy these regions (Figure 1F). These results establish that p53 can activate RFX7 to employ the RFX7 GRN revealing a novel p53-RFX7 signaling axis.

**Figure 1.**
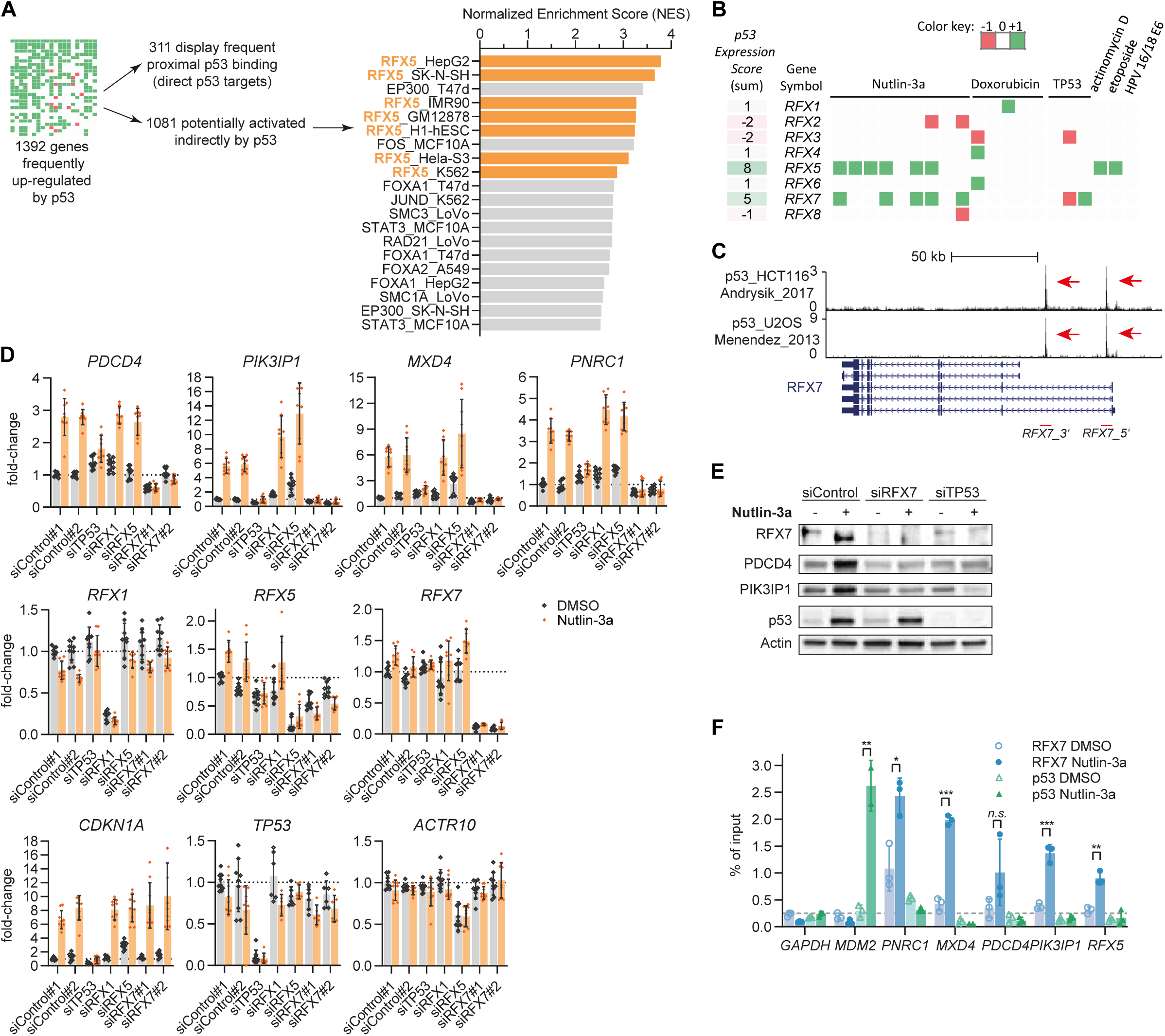
The p53 target RFX7 mediates p53-dependent gene activation. **(A)** Our previous meta-analysis identified 1392 genes as frequently up-regulated by p53 (*p53 Expression Score* ≥ 5). Out of these genes, 311 displayed frequent p53 binding within 2.5kb of their TSS and represent high-probability direct p53 targets (20). Transcription factors enriched for binding within 500 bp upstream from the TSS of the remaining 1081 genes were identified using publicly available ChIP-seq data. ChIP-seq datasets with a normalized enrichment score (NES) > 2.5 are displayed. **(B)** The p53-dependent regulation of RFX family encoding genes across 20 datasets from a meta-analysis (20). Genes were identified as significantly up-regulated (green; +1), down-regulated (red; -1), or not significantly differentially regulated (white; 0). The *p53 Expression Score* represents the summary across all 20 datasets. **(C)** Genome browser snapshot displaying publicly available p53 binding signals from Nutlin-3a treated U2OS and HCT116 cells at the *RFX7* gene locus. Red arrows indicate two p53 binding signals in *RFX7* intron1, one located 5’ (*RFX7_5’*) and one 3’ (*RFX7_3’*). **(D)** RT-qPCR data of selected direct RFX7 targets in U2OS cells. Normalized to siControl#1 DMSO. *ACTR10* served as negative control. *TP53*, *RFX1*, *RFX5*, and *RFX7* are shown as knockdown controls. *CDKN1A* is a positive control for p53 induction by Nutlin-3a. Mean and standard deviation is displayed. Statistical significance obtained through a two-sided unpaired t-test, n = 9 replicates (3 biological with 3 technical each). **(E)** Western blot analysis of RFX7, p53, PDCD4, PIK3IP1, and actin (loading control) levels in U2OS cells transfected with siControl, siRFX7, or siTP53 and treated with Nutlin-3a or dimethyl sulfoxide (DMSO) solvent control. **(F)** RFX7 and p53 ChIP-qPCR of selected RFX7 targets in Nutlin-3a and DMSO control-treated U2OS cells. *GAPDH* served as negative control. *MDM2* served as positive control for p53 binding. Mean and standard deviation is displayed. Statistical significance obtained through a two-sided unpaired t-test, n = 3 technical replicates.

To test whether the p53-dependent function of RFX7 is cell type-specific or represents a more ubiquitous mechanism, we employed the colorectal cancer cell line HCT116 and the hTERT-immortalized non-cancerous retina pigmented epithelium cell line RPE-1, both of which possess wild-type p53. ChIP-qPCR analysis confirmed that p53 binds to two sites in RFX7 intron1 in U2OS, HCT116, and RPE-1 cells (Figure 2A). Similar to our results from U2OS cells (Figure 1D and E), *PDCD4*, *PIK3IP1*, *MXD4*, and *PNRC1* were induced upon Nutlin-3a treatment in HCT116 and RPE-1 cells in a p53 and RFX7-dependent manner, while *CDKN1A* was not affected by RFX7 depletion (Figure 2B). Protein levels of PDCD4 and PIK3IP1 largely followed the p53-RFX7-dependent regulation of their mRNAs (Figure 2C). Given the diversity of the investigated cell lines, our data suggest that the novel p53-RFX7 signaling axis influences numerous cell types. Together, our findings establish RFX7 as a novel direct p53 target that extends p53-dependent gene activation to potent tumor suppressor genes in numerous cell types.

**Figure 2.**
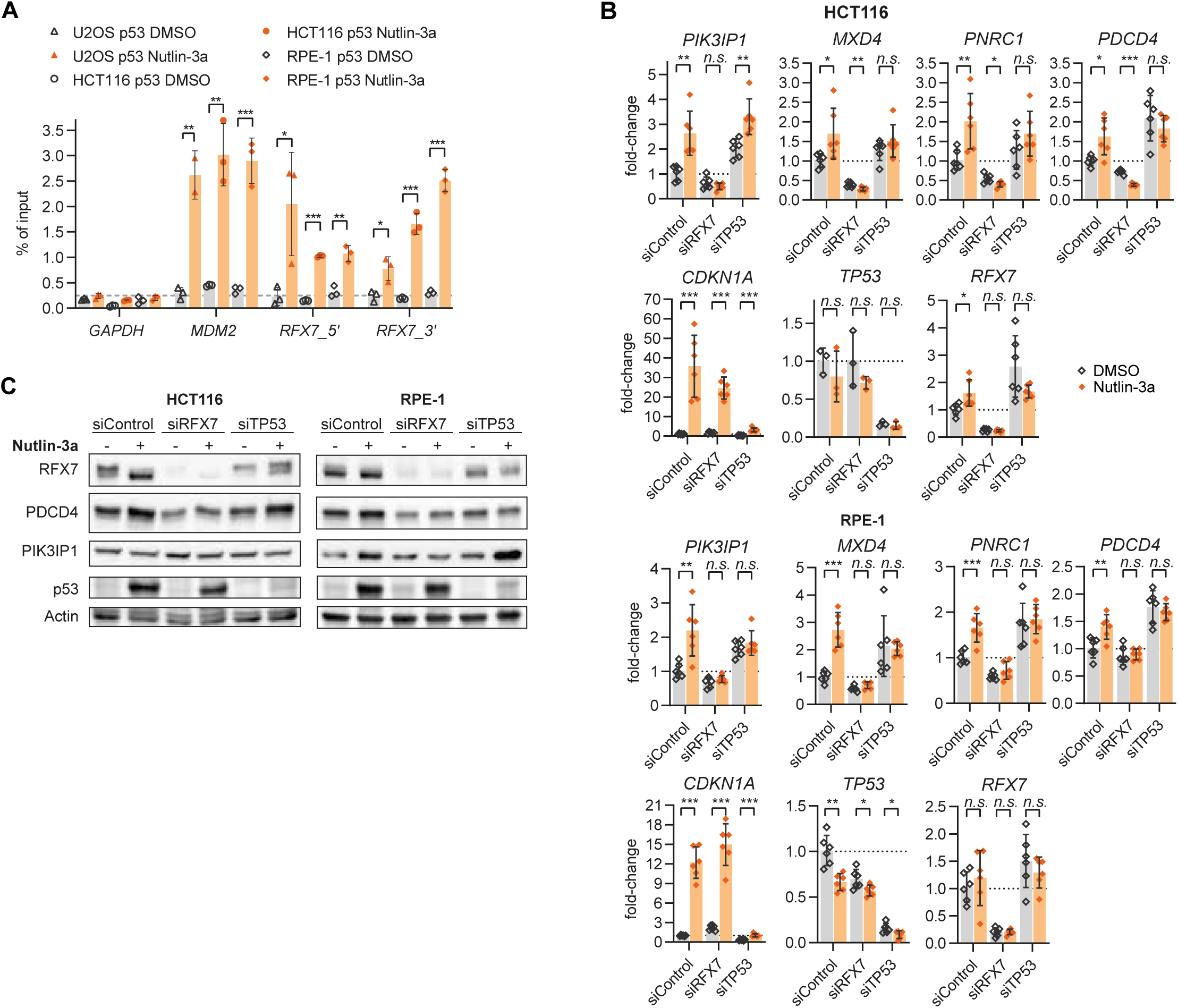
The direct p53 target RFX7 functions in numerous cell types. **(A)** ChIP-qPCR of p53 binding to *GAPDH* (negative control), *MDM2* (positive control), and the 5’ (*RFX7*_5’) and 3’ (*RFX7*_3’) sites in *RFX7* intron1 from U2OS, HCT116, and RPE-1 cells treated with Nutlin-3a or DMSO solvent control. Statistical significance obtained through a two-sided unpaired t-test, n = 3 technical replicates. **(B)** RT-qPCR data of *PDCD4*, *PIK3IP1*, *MXD4*, and *PNRC1* in HCT116 and RPE-1 cells. Normalized to *ACTR10* negative control and siControl DMSO sample. Mean and standard deviation is displayed. Statistical significance obtained through a two-sided unpaired t-test, n = 6 replicates (2 biological with 3 technical each). *TP53* and *RFX7* are shown as knockdown controls. *CDKN1A* is a positive control for p53 induction by Nutlin-3a. **(C)** Western blot analysis of RFX7, p53, PDCD4, PIK3IP1, and actin (loading control) levels in HCT116 and RPE-1 cells transfected with siControl, siRFX7, or siTP53 and treated with Nutlin-3a or DMSO solvent control.

### The RFX7 DNA binding landscape enriches proximal promoter regions

The identification of p53 as an upstream regulator of RFX7 enabled us to induce RFX7 levels and activity pharmacologically. Although RFX7 emerged as a potent suppressor of lymphoid cancers and putative cancer driver in Burkitt lymphoma (4, 5, 9, 62), the mechanisms underlying its tumor suppressor function remain elusive. Given that RFX7 is a transcription factor, it seems natural that its tumor suppressor function is mediated through its target genes. To identify RFX7 target genes genome-wide, we performed ChIP-seq in Nutlin-3a and DMSO control-treated U2OS, HCT116, and RPE-1 cells (Figure 3A, Supplementary Table 1). Substantially more RFX7 binding sites were identified in Nutlin-3a compared to DMSO control-treated cells (Supplementary Table 1), underlining the importance of inducing RFX7 levels and activity to identify RFX7-dependent genome regulation. We focused further investigations on sites occupied by RFX7 across all three cell types upon Nutlin-3a treatment (Figure 3A). RFX7 binding sites are phylogenetically conserved (Figure 3B) and predominantly located near transcriptional start sites (TSSs) (Figure 3C). *De novo* search for motifs underlying RFX7 binding sites revealed an X-box that is commonly recognized by the RFX family (1) and a CCAAT-box known to recruit NF-Y (63) (Figure 3D). Corroborating the ChIP-qPCR results (Figure 1F), Nutlin-3a treatment led to increased RFX7 DNA occupancy genome-wide (Figure 3E). For example, the p53-RFX7-regulated genes *PNRC1* and *MXD4* (Figure 1D and E) display RFX7 binding near their TSSs, which increased upon Nutlin-3a treatment (Figure 3F). Enrichment analysis identified RFX5 and its co-factor CIITA, FOS, NF-Y, CREB1, EP300, and STAT3, among others, to share a significant number of binding sites with RFX7 (Figure 3G), which indicates that RFX5 and RFX7 bind to similar sites and that RFX7, similar to RFX5 (64), may cooperate with the CCAAT-box binding NF-Y. Given that RFX5 but not RFX1 was identified to share binding sites with RFX7 (Figure 3G), we compared the RFX7 X-box motif (Figure 3D) with known X-box motifs of the RFX family to identify potential differences. RFX7 shares the X-box motif with other RFX family members, but it shows a clear distinction. While RFX1-3 bind to a palindromic X-box comprising two half-sites (1), RFX7 – similar to RFX5 – binds to an X-box with only one half-site (Figure 3H). Although the RFX family shares a conserved DBD, there are differences in their motif recognition, which offers an explanation for sites that are exclusively bound by RFX7 and RFX5. However, comparing the binding site repertoire of RFX5 and RFX7 revealed a substantial difference as RFX7 occupies only a small subset of RFX5 binding sites (Figure 3I). Together, these findings show that RFX7 differs markedly from all other members of the RFX transcription factor family, including its phylogenetically closest sibling RFX5.

**Figure 3.**
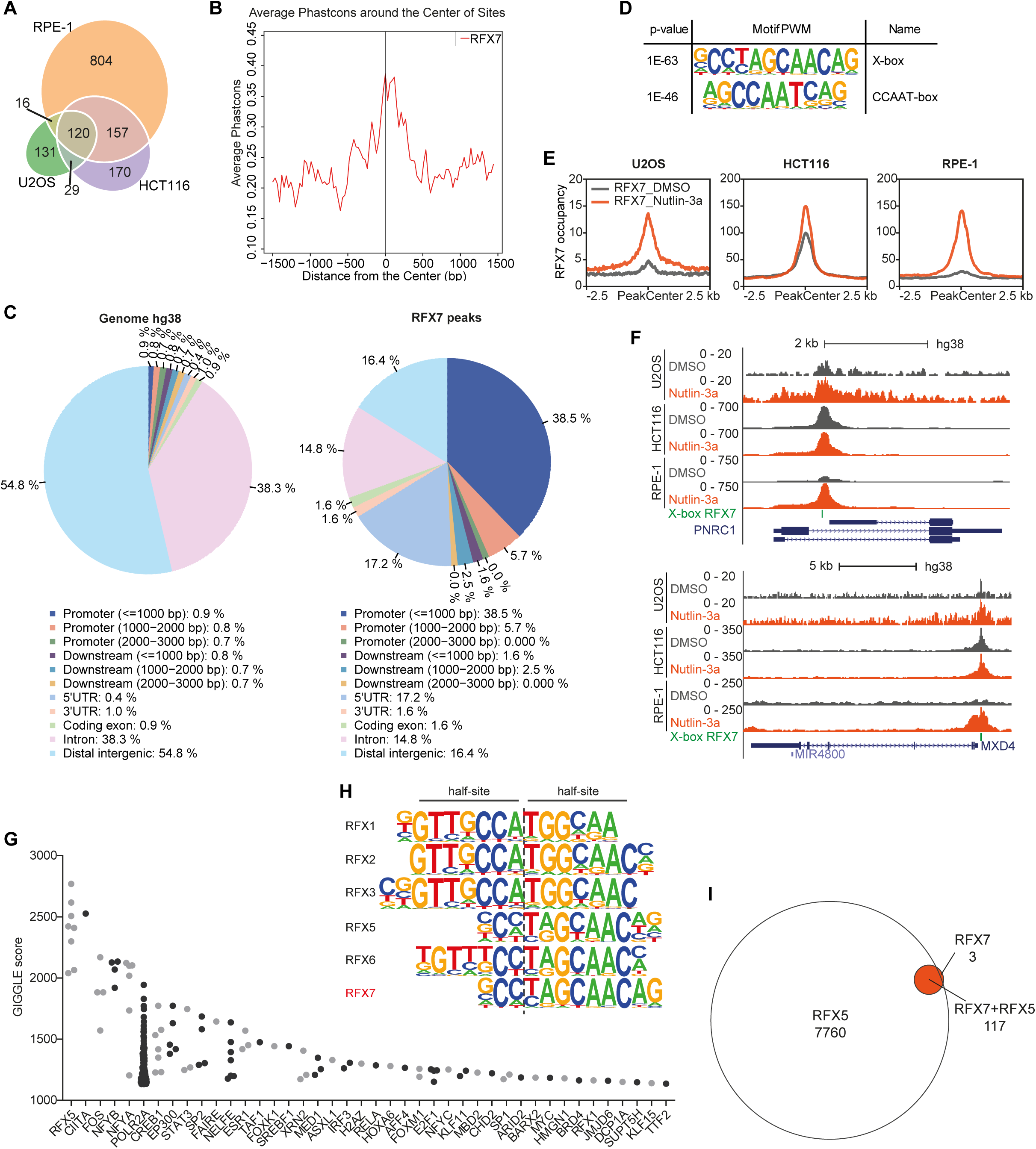
The DNA binding landscape of RFX7. **(A)** Number of RFX7 ChIP-seq peaks identified in Nutlin-3a-treated U2OS, HCT116, and RPE-1 cells. **(B)** Average vertebrate PhastCons conservation score at the 120 common RFX7 binding sites. **(C)** CEAS Enrichment on annotation analysis (38) for the 120 common RFX7 peaks compared to the human genome hg38. **(D)** Top motifs identified by *de novo* motif analysis using HOMER under the 120 peaks commonly identified in all three cell lines. **(E)** Mean RFX7 occupancy (ChIP-seq read counts) at the 120 common RFX7 binding sites. **(F)** Genome browser images displaying RFX7 ChIP-seq signals and predicted X-boxes at the *PNRC1* and *MXD4* gene loci. **(G)** Transcription factor ChIP-seq peak sets from CistromeDB that overlap significantly with the 120 common RFX7 binding sites. **(H)** Comparison of known X-box motifs from RFX family members with the X-box we identified for RFX7. Known motifs were obtained from the HOMER motif database. **(I)** The overlap of the 120 common RFX7 binding sites with 7877 RFX5 binding sites supported by at least 5 out of 10 ChIP-seq datasets.

### RFX7 functions as a *trans*-activator to alter the transcriptome

To complement the RFX7 DNA binding landscape, we identified the RFX7-regulated transcriptome through RNA-seq analyses of U2OS, HCT116, and RPE-1 cells treated with Nutlin-3a or DMSO control. RNA-seq data confirmed significant Nutlin-3a-induced up-regulation of *RFX5* and *RFX7*, while *RFX1* is not induced (Figure 4A). Depletion of RFX7 caused up and down-regulation of several hundred genes (Figure 4B, Supplementary Table 2). While RFX7-dependent regulation was observed to be cell line-specific at large, we identified multiple genes affected by RFX7 depletion across cell lines. Genes down-regulated upon RFX7 knockdown enriched for RFX5 binding and NF-Y motifs. In contrast, up-regulated genes enriched for AP-1 (JUN/FOS) binding and motifs (Figure 4C). The fact that RFX7 occupies similar sites as RFX5 and NF-Y (Figure 3G) already indicates that RFX7 may predominantly *trans*-activate its target genes. Indeed, integration of ChIP-seq and transcriptome data corroborates RFX7’s *trans*-activator function (Supplementary Figure 1B). In turn, the set of genes directly activated by RFX7 might indirectly convey repressive effects on the highly cell-type specific AP-1 signaling.

**Figure 4.**
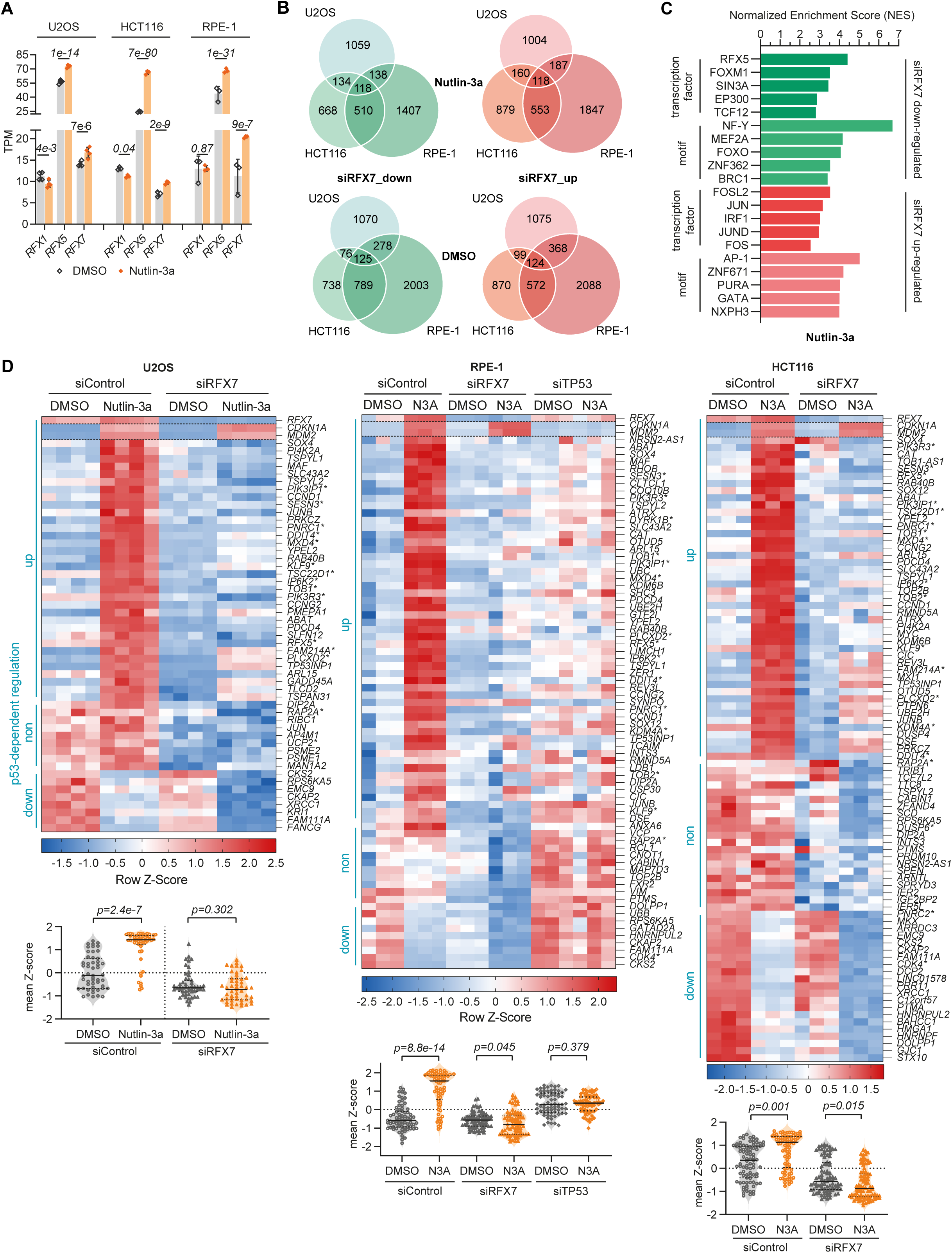
The RFX7-regulated transcriptome. **(A)** TPM (Transcripts Per Kilobase Million) expression values of *RFX1*, *RFX5*, and *RFX7* obtained from RNA-seq analysis from U2OS, HCT116, and RPE-1 cells treated with Nutlin-3a or DMSO solvent control. Statistical significance data from DESeq2 analysis. **(B)** Number of genes significantly (FDR < 0.01) up (log_2_FC ≥ 0.25; red venn diagrams) or down-regulated (log_2_FC ≤ 0.25; green venn diagrams) following siRFX7 treatment in DMSO (bottom venn diagrams) and Nutlin-3a-treated (upper venn diagrams) U2OS, HCT116, and RPE-1 cells. **(C)** Top 5 transcription factors and binding motifs enriched among genes significantly up or down-regulated following RFX7 depletion in at least two Nutlin-3a-treated cell lines. **(D)** Heatmap of RNA-seq data for direct RFX7 target genes that bind RFX7 within 5 kb from their TSS according to ChIP-seq data and are significantly (FDR < 0.01) down-regulated (log_2_FC ≤ -0.25) following RFX7 depletion in Nutlin-3a treated U2OS, HCT116, and RPE-1 cells. Significant (FDR < 0.01) p53-dependent regulation is indicated at the left. Asterisks (*) indicate conserved Rfx7-dependent expression in mouse spleen or bone marrow (17) (Supplementary Table 3). Violin plots correspond to the heatmaps and display the mean Z-score of all these genes for the different treatment conditions. The median is indicated by a black line. Statistical significance obtained using a two-sided paired t-test.

### The RFX7 target gene network comprises multiple tumor suppressors and responds to stress

We integrated the RFX7 DNA binding landscape and the RFX7-regulated transcriptome to infer potential direct RFX7 target genes genome-wide. We identified 51, 87, and 73 potential direct RFX7 targets in U2OS, HCT116, and RPE-1 cells, respectively, and these direct RFX7 targets include *PDCD4*, *PIK3IP1*, *MXD4*, and *PNRC1* (Figure 4D). Most strikingly, target genes up-regulated through the p53-RFX7 axis comprise additional tumor suppressor genes, such as *ABAT* (65), *CCNG2* (66), *IP6K2* (67), *OTUD5* (68), *REV3L* (69), *RPS6KA5* (also known as *MSK1*) (70), *TOB1* (71), *TSC22D1* (72), and *TSPYL2* (73). Most direct RFX7 targets were up-regulated in response to Nutlin-3a treatment in siControl-transfected cells, and the up-regulation was impaired or abrogated when RFX7 was depleted (Figure 4D). Notably, 15, 19, and 16 (20-30%) of direct RFX7 target genes identified in U2OS, HCT116, and RPE-1 cell, respectively, displayed conserved Rfx7-dependent expression in mouse spleen or bone marrow (Supplementary Table 3) (17). Direct RFX7 target genes down-regulated upon Nutlin-3a treatment comprise cell cycle genes, including *DOLPP1*, *XRCC1*, *CDK4*, *CKAP2*, *FAM111A*, and *CKS2*, that become down-regulated through the *trans*-repressor complex DREAM (20). These Nutlin-3a-repressed genes display a more marked decrease in mRNA levels when RFX7 is missing, suggesting that RFX7 partially counteracts and limits their p53-dependent down-regulation. Given that RPE-1 is no established cell line model in p53 research, we provide data showing that depletion of p53 in RPE-1 abrogated the Nutlin-3a-induced regulation of all those genes (Figure 4D). Direct RFX7 target genes identified in at least two of the three cell lines comprise a set of 57 genes (Table 1). In addition to regulating multiple tumor suppressors directly, our data reveal a large p53-dependent subnetwork co-directed by RFX7 (Supplementary Figure 2, Supplementary Table 2), further highlighting the impact of the novel p53-RFX7 signaling axis.

**Table 1.** Direct RFX7 target genes. Set of 57 direct RFX7 target genes identified as bound by RFX7 and down-regulated upon RFX7 knockdown in at least two of the three cell lines (Figure 4D). Genes within the 19-gene-set used for survival analyses are marked bold.

|  |  |  |  |  |
| --- | --- | --- | --- | --- |
| **ABAT** | *DIP2A* | *KLF9* | *PTMS* | *TOB2* |
| *ARL15* | *DOLPP1* | *MAF* | *RAB40B* | *TOP2B* |
| *ATRX* | *DSE* | ***MXD4*** | *RAP2A* | ***TP53INP1*** |
| *CABIN1* | *EMC9* | *NRSN2-AS1* | *REV3L* | *TSC22D1* |
| *CAT* | *FAM111A* | *OTUD5* | ***RFX5*** | ***TSPYL1*** |
| ***CCND1*** | *FAM214A* | ***PDCD4*** | *RMND5A* | ***TSPYL2*** |
| ***CCNG2*** | *HNRNPUL2* | *PI4K2A* | *RPS6KA5* | *UBE2H* |
| *CDK4* | *INTS3* | ***PIK3IP1*** | ***SESN3*** | *XRCC1* |
| *CIC* | ***IP6K2*** | ***PIK3R3*** | ***SLC43A2*** | ***YPEL2*** |
| *CKAP2* | ***JUNB*** | ***PLCXD2*** | *SOX12* |  |
| *CKS2* | *KDM4A* | ***PNRC1*** | *SOX4* |  |
| *DDIT4* | *KDM6B* | *PRKCZ* | ***TOB1*** |  |

Integration of our meta-analysis data (20) showed that most direct RFX7 targets become up-regulated by p53 in various cell types and in response to multiple stimuli (Figure 5A). Consequently, we tested whether RFX7 affected their regulation in response to cellular stress. To this end, we employed Doxorubicin and Actinomycin D, which are well-established to induce the p53 program (20). Doxorubicin is a topoisomerase II inhibitor that causes DNA double-strand breaks while Actinomycin D inhibits rRNA transcription inducing ribosomal stress. *PDCD4*, *PIK3IP1*, *MXD4*, and *PNRC1* were up-regulated in response to Nutlin-3a, Actinomycin D, and Doxorubicin treatment. The up-regulation was attenuated when p53 or RFX7 were depleted. The direct p53 target *CDKN1A* was up-regulated p53-dependent and RFX7-independent (Figure 5B). These results identify RFX7 as a missing link to up-regulate numerous tumor suppressor genes in response to stress.

**Figure 5.**
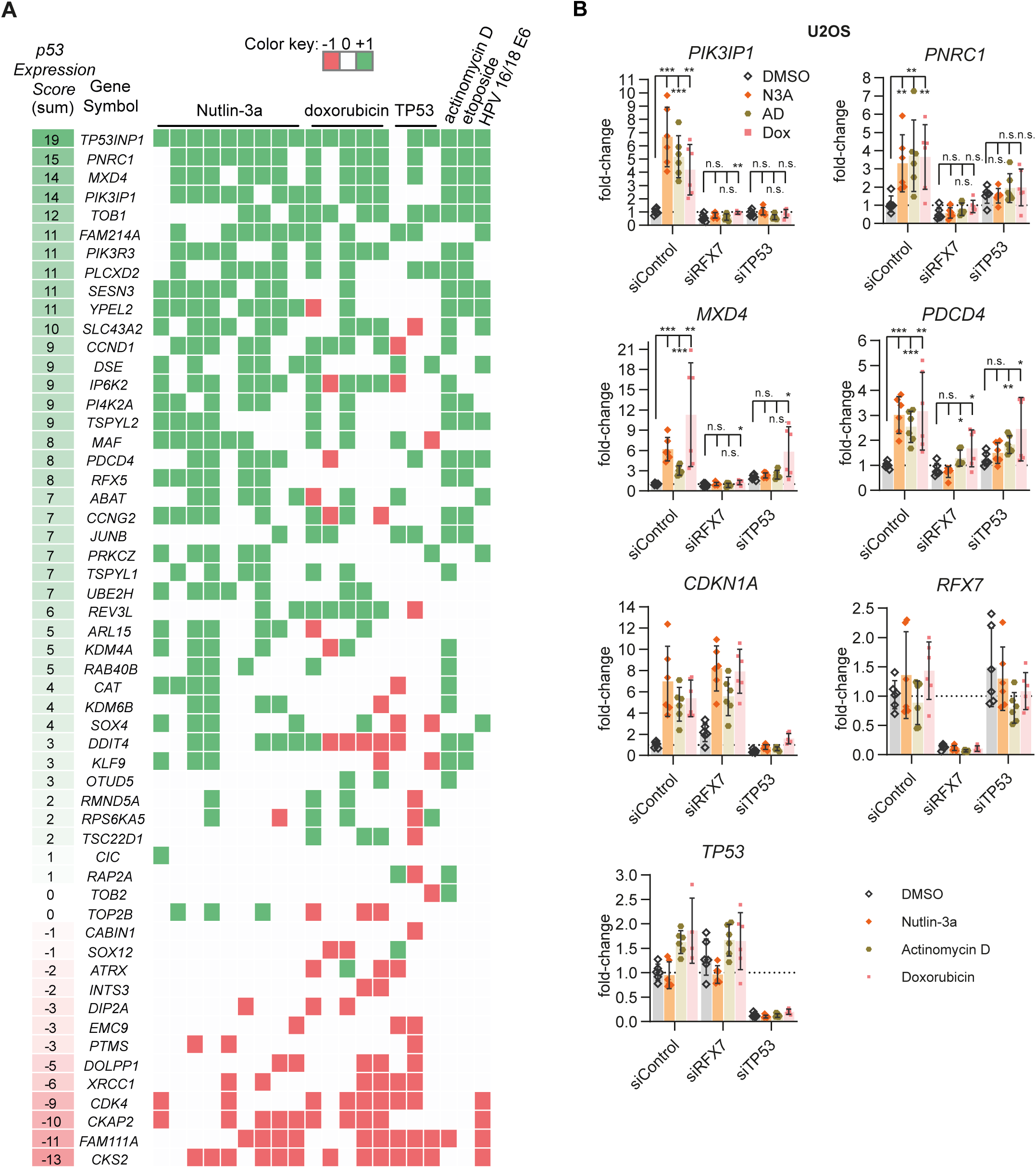
RFX7 up-regulates its target genes in response to stress. **(A)** The p53-dependent regulation of direct RFX7 target genes (Table 1) across 20 datasets from a meta-analysis (20). Genes were identified as significantly up-regulated (green; +1), down-regulated (red; -1), or not significantly differentially regulated (white; 0). The *p53 Expression Score* represents the summary across all 20 datasets. No meta-analysis data was available for *NRSN2-AS1* and *HNRNPUL2*. **(B)** RT-qPCR data from U2OS cells depleted for RFX7 or p53 and treated with DMSO control, Nutlin-3a (N3A), Actinomycin D (AD), and Doxorubicin (Dox). RT-qPCR data normalized to *ACTR10* and siControl DMSO levels. Mean and standard deviation is displayed. Statistical significance of RT-qPCR data obtained through a two-sided unpaired t-test, n = 6 replicates (2 biological with 3 technical each).

### High RFX7 target gene expression is associated with better prognosis in cancer patients and cell differentiation

We and others identified RFX7 as a putative cancer driver in Burkitt Lymphoma (4, 5), and mouse data confirmed its tumor suppressor function in lymphoma development (9). Here, we identified RFX7 to induce well-established tumor suppressor genes in numerous cell types (Figure 4D). To assess whether RFX7 may affect cell growth and tumor development also in cell types outside the lymphoid lineage, we resorted to publicly available cell viability data from the DepMap project (51). Intriguingly, *RFX7* knockdown increased the viability of the majority of 343 cell lines tested, while the viability of lymphoma cell lines increased the most (Figure 6A). Given that RFX7 appears to restrict cell growth across a wide range of cell types, we sought to assess the potential role of RFX7 signaling in numerous cancer types. Therefore, we resorted to the cancer genome atlas (TCGA) that comprises patient data from 33 cancer types (74), and we tested whether RFX7 target gene expression is associated with patient survival. To avoid confounding effects from cell cycle genes, which are well-established to be associated with worse prognosis across cancer types (75), we used a subset of 19 direct RFX7 target genes that are frequently up-regulated by p53. Strikingly, higher expression of these direct RFX7 targets correlates significantly with better prognosis across the whole TCGA pan-cancer cohort (Figure 6B and Supplementary Figure 3). Survival analyses using data from the 33 individual cancer types revealed that in 11 out of the 33 individual cancer types higher expression of the RFX7 targets correlates significantly with better prognosis (Figure 7). These findings indicate that RFX7 signaling is frequently de-regulated in cancer. Together, these data indicate a ubiquitous role of RFX7 in restricting cell growth and potential clinical implications of this new signaling axis in numerous cancer types.

**Figure 6.**
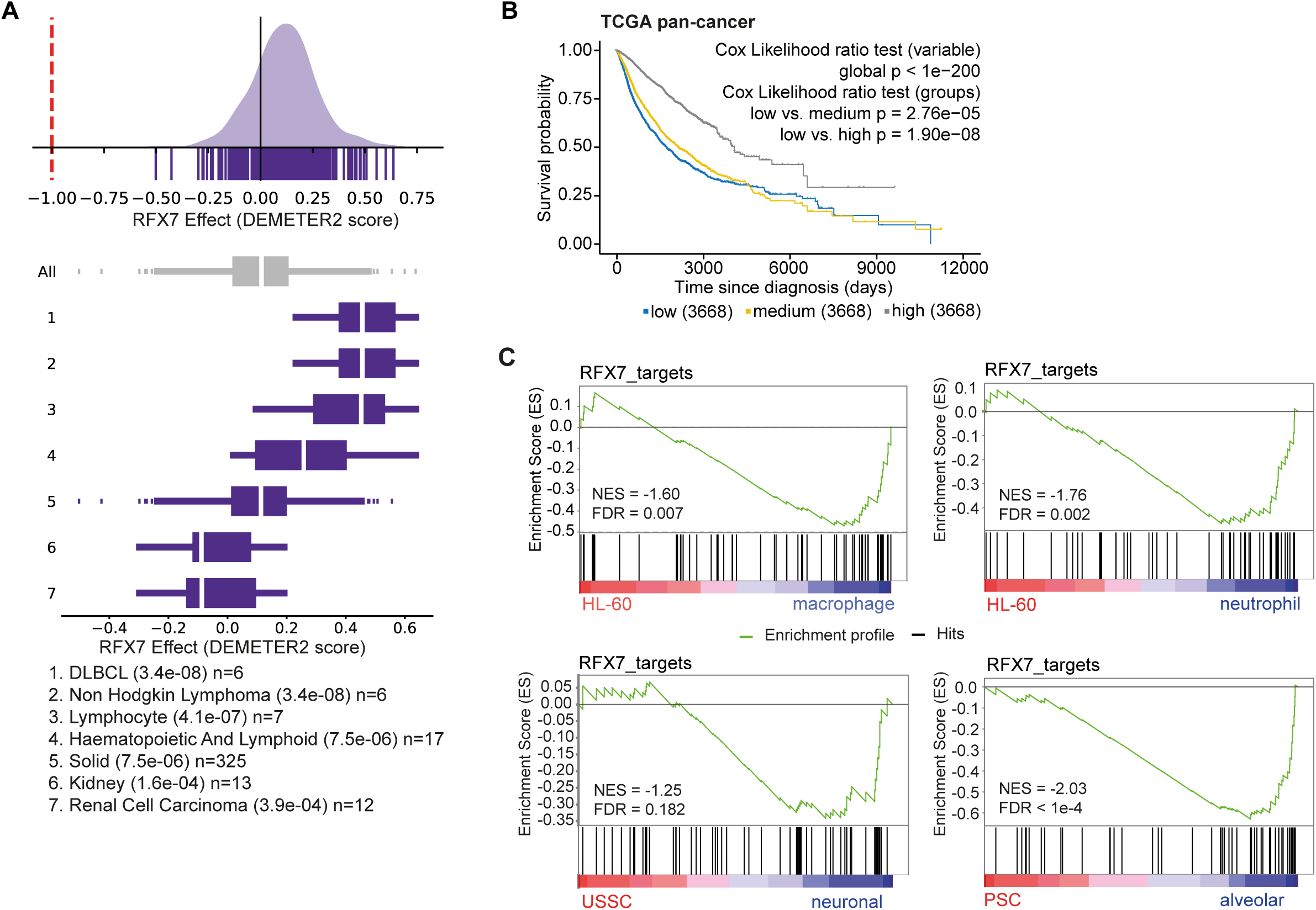
RFX7 limits cell viability, and RFX7 target gene expression correlates with good prognosis in cancer and cell differentiation. **(A)** Cell viability data from depmap.org (51). DEMETER2 dependency scores (52) are based on RNAi mediated knockdown of RFX7 in 343 cell lines. Top panel displays data for all 343 cell lines and bottom panel displays groups of cell lines that show DEMETER2 scores significantly different to all other cell lines. Groups are based on tissue origin. Negative dependency scores reflect decreased cell viability upon loss of the target gene, while positive scores indicate increased cell viability. **(B)** Kaplan-Meier plot of patients from the TCGA pan-cancer cohort. Patients were grouped into low, medium, and high based on the rank expression of 19 direct RFX7 target genes that display a *p53 Expression Score* > 5 (20). Statistical significance obtained through the Cox proportional hazards (PH) model (Cox likelihood ratio test variable). To correct for major confounding factors, cancer type, gender, and age were included into the multivariate regression analysis. Statistical significance of the rates of occurrence of events over time between the groups was obtained using the fitted Cox PH model (Cox likelihood ratio test groups). Complementary data displayed in Supplementary Figure 3. **(C)** Gene set enrichment analysis (GSEA) of 57 direct RFX7 target genes in human p53-negative HL-60 promyelocytes differentiated into macrophages or neutrophils (upper panels) (47), human umbilical cord blood-derived unrestricted somatic stem cells (USSC) differentiated into neuronal-like cells (48) and human pluripotent stem cells (PSC) differentiated into lung alveolar cells (bottom panels) (49).

**Figure 7.**
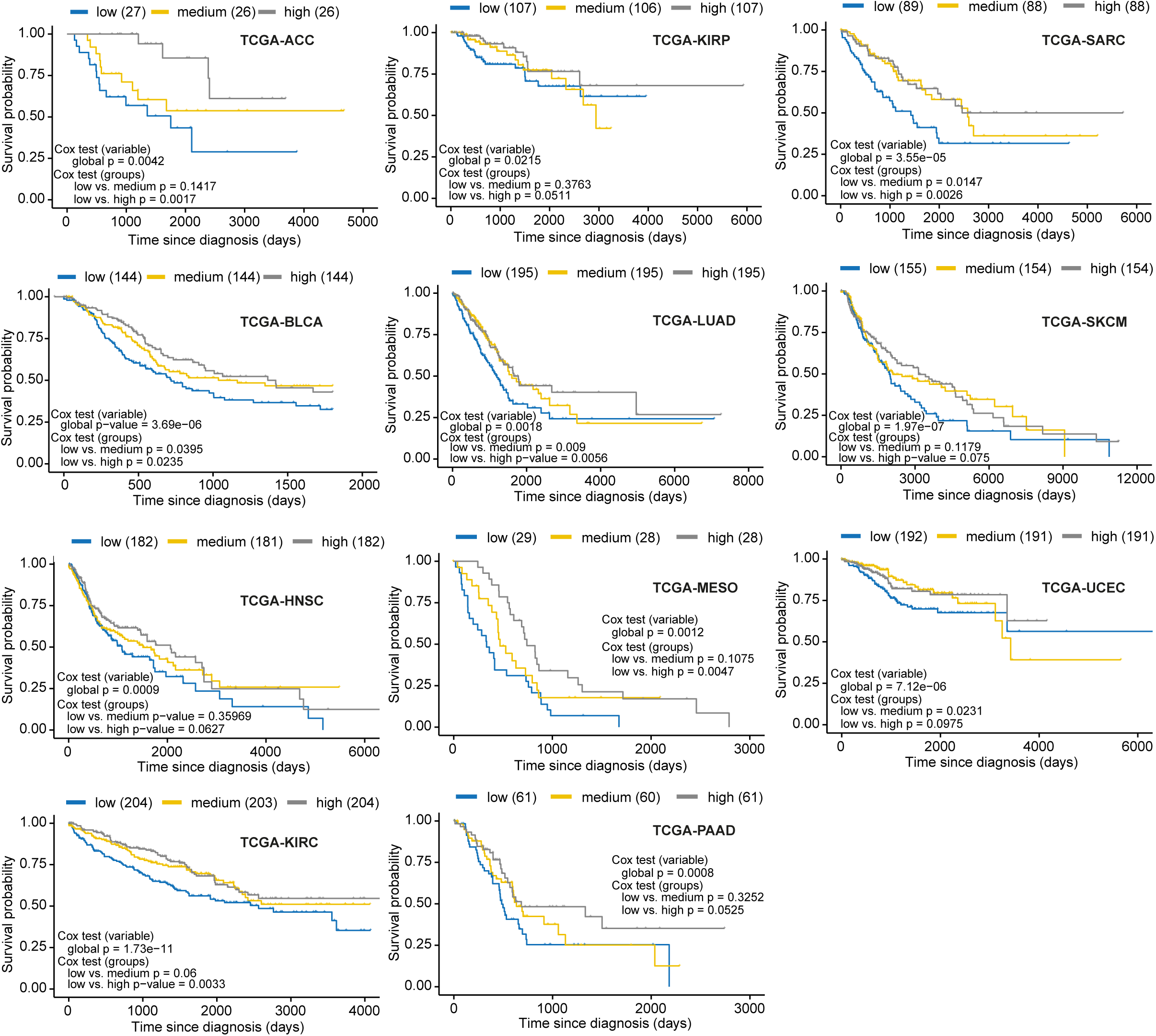
RFX7 target gene expression correlates significantly positive with good prognosis in 11 cancer types. Kaplan-Meier plots of patients from TCGA cohorts. Patients were grouped into low, medium, and high based on the rank expression of 19 direct RFX7 target genes that display a *p53 Expression Score* > 5 (20). 11 out of 33 cancer types are displayed that show a significantly (Cox likelihood ratio test variable < 0.05 and Cox likelihood ratio test group low vs high < 0.1) better prognosis when the expression of the RFX7 targets is higher. Only one cancer type (TCGA-COAD) showed a significantly poorer prognosis. Statistical significance obtained through the Cox proportional hazards (PH) model (Cox test variable). To correct for major confounding factors, gender and age were included into the multivariate regression analysis. Statistical significance of the rates of occurrence of events over time between the groups was obtained using the fitted Cox PH model (Cox test groups).

Cell differentiation represents an anti-proliferative mechanism that is typically circumvented by cancer (76). More differentiated cancer cells are characterized as low grade and are often associated with a favorable prognosis. Notably, RFX7 orthologs have been shown to play a role in the development of murine natural killer cells (17) and in the neural development of frogs (16). Given that several direct RFX7 targets have been associated with cell differentiation, such as the MYC antagonist MXD4 (also known as MAD4) (77), we tested whether RFX7 could play a more general role during differentiation. Therefore, we assessed the expression of RFX7 target genes when human p53-negative HL-60 promyelocytes differentiated into macrophages or neutrophils. Interestingly, RFX7 target gene expression correlated significantly positively with macrophage and neutrophil differentiation (Figure 6C), indicating a potential role for RFX7 in hematopoietic differentiation that is independent of p53. Further, RFX7 target gene expression correlated positively with the differentiation of human umbilical cord blood-derived unrestricted somatic stem cells into neuronal-like cells (Figure 6C), which is in agreement with the reported role of RFX7 in the neural development of frogs (16) and its association with neurological diseases (14, 15). Intriguingly, the expression of RFX7 target genes correlates significantly positively also with the differentiation of human pluripotent stem cells into lung alveolar cells (Figure 6C). Together, these results indicate a potentially widespread role of RFX7 in promoting cell differentiation.

### RFX7 sensitizes cells to Doxorubicin and promotes apoptosis

The potentially widespread role of RFX7 in cancer (Figure 6B and 7) and its activation in response to cellular stress (Figure 5) prompted us to investigate the role of RFX7 in the stress response. To this end, we challenged U2OS osteosarcoma and HCT116 colorectal cancer cells with different concentrations of Doxorubicin. Intriguingly, WST-1 assays showed that RFX7 depletion significantly increased the viability of U2OS and HCT116 cells challenged with low concentrations of Doxorubicin, with HCT116 showing the highest benefit (Figure 8A). Confirming previous results (78), depletion of p53 did not increase the viability. We further assessed the response in HCT116 cells, and validated increased viability in response to Doxorubicin through clonogenic and Annexin V assays (Figure 8B and C). Importantly, Annexin V assays revealed that the increased cell viability upon RFX7 depletion was associated with significantly reduced apoptosis (Figure 8C). Thus, RFX7 appears to sensitize cells to Doxorubicin through promoting apoptosis, indicating a role of RFX7 in cell fate determination in response to stress.

**Figure 8.**
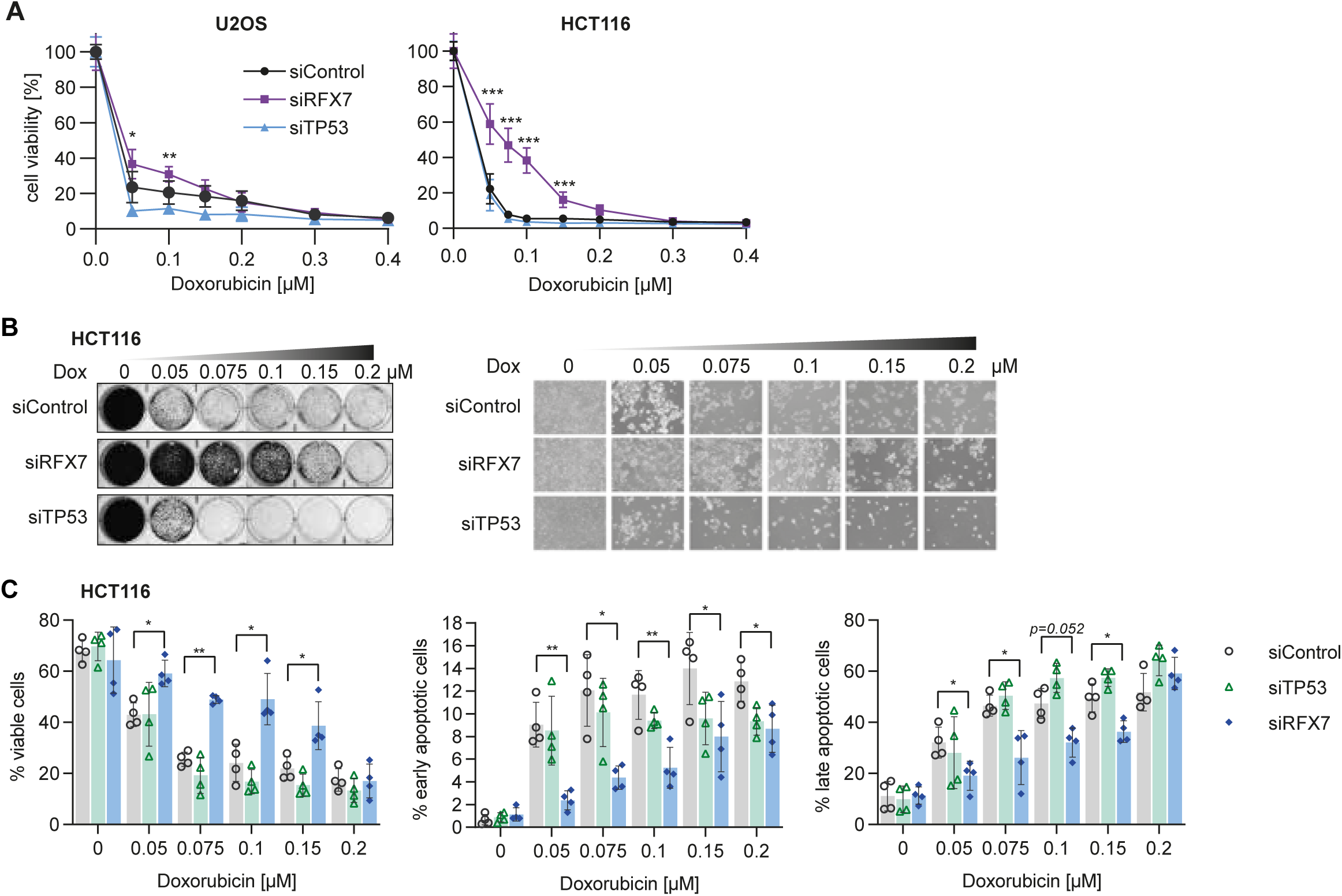
RFX7 sensitizes to Doxorubicin through promoting apoptosis. **(A)** WST-1 assay of U2OS and HCT116 cells challenged with different concentrations of Doxorubicin. Mean and standard deviation is displayed. Statistical significance between siControl and siRFX7 obtained through a Sidak-corrected two-way ANOVA test, n = 11 (U2OS) or 9 (HCT116) biological replicates. **(B)** Clonogenic assay of HCT116 cells challenged with different concentrations of Doxorubicin (left) and complementary brightfield images (right). **(C)** Annexin V assay of HCT116 cells transfected with siControl, siTP53, or siRFX7 and treated with indicated concentrations of Doxorubicin. Viable cells (negative for Annexin V and PI), early apoptotic cells (positive for Annexin V, negative for PI), and late apoptotic cells (positive for Annexin V and PI) were quantified through flow cytometry. Relative numbers of 50,000 cells from n = 4 biological replicates are displayed. Mean and standard deviation is displayed. Statistical significance between siControl and siRFX7 obtained through a two-sided paired t-test.

## Discussion

p53 is the best-known tumor suppressor, but it remains unclear how it regulates large parts of its GRN. Our findings place the understudied transcription factor RFX7 immediately downstream of p53 in regulating multiple genes. RFX7 emerged recently as an essential regulator of lymphoid cell maturation (17) and a putative cancer driver mutated in hematopoietic neoplasms (62). While these observations are in agreement with the maximal expression of *RFX7* in lymphoid tissue (3, 17), our results using human osteosarcoma, colorectal cancer, and non-cancerous retinal pigment epithelial cells establish a ubiquitous role of RFX7 in regulating known tumor suppressors and in serving as a crucial regulatory arm of the p53 tumor suppressor. We establish p53 as the first regulator of the novel tumor suppressor RFX7 and exploit this regulatory connection to chart RFX7’s target gene network in three distinct cell systems. Most importantly, the RFX7 network comprises multiple established tumor suppressor offering an explanation for RFX7’s tumor suppressor role. For example, similar to the lymphoma-promoting loss of Rfx7 in a mouse model (9), mice carrying a knockout of the RFX7 targets PDCD4 and REV3L displayed spontaneous lymphomagenesis (57, 69). Similar to the transcription factor p53, RFX7 appears to orchestrate its tumor suppressive function through multiple target genes.

The general importance of RFX7 signaling in cancer biology is exemplified by the better prognosis of patients with medium to high expression of RFX7 targets across the pan-cancer cohort (Figure 6B). While frequent mutations in *RFX7* so far have been identified only in Burkitt lymphoma (4, 5, 62), the altered expression of direct RFX7 target genes across numerous cancer types (Figure 6B and 7) indicates that RFX7 signaling is recurrently de-regulated in cancer. High expression of RFX7 target genes during differentiation (Figure 6C) and RFX7’s apoptosis-promoting function in response to stress (Figure 8) indicate a widespread role of RFX7 in cell fate determination and may at least in part account for the better prognosis observed in cancer patients with higher RFX7 target gene expression. RFX7 promoting apoptosis in response to Doxorubicin treatment may be attributed to its target IP6K2, an established inducer of apoptosis (67).

The direct RFX7 target genes *RPS6KA5* (*MSK1*) and *ARL15* (Figure 4D) may explain the link between *RFX7* alteration and increased waist-hip-ratio (13), as both genes have been associated with obesity and high waist-hip-ratio (79). Furthermore, RFX7 directly regulates multiple transcription factors, including JUNB, KLF9, MAF, MXD4, RFX5, SOX4, SOX12, and TSC22D1, as well as chromatin modifiers, which may affect many RFX7-regulated genes that are not bound by RFX7 itself (Figure 4D, Supplementary Figure 2).

In summary, our findings suggest that the RFX7 signaling pathway represents a novel growth regulatory mechanism that is activated in response to stress and p53. Given the importance of the discovered regulatory connection, we expect our data to be essential in triggering further research into RFX7’s regulatory network, potentially leading to new diagnostic and therapeutic approaches.

## Acknowledgements

This work was supported by the German Research Foundation (DFG) [research grant FI 1993/2-1 to M.F.] and the German Federal Ministry for Education and Research (BMBF) [031L016D to S.H.; ICGC MMML-Seq 01KU1002A-J and ICGC-Data Mining 01KU1505-C and G to R.S. and S.H.]. The FLI is a member of the Leibniz Association and is financially supported by the Federal Government of Germany and the State of Thuringia.

We gratefully acknowledge the FLI Core Facilities DNA-sequencing (Ivonne Görlich and Cornelia Luge) and Flow Cytometry (Michelle Burkhardt) for their technical support. We thank Bernhard Schlott for the kind gift of p53 antibody.

## Author Contributions

M.F. conceived the study. M.F. and S.H. supervised the work. M.F., L.C., and S.H. designed the experiments. L.C., K.S., M.F., S.F., D.H., and L.S. performed the experiments. K.R., M.F., S.H.B., and S.H. performed the computational analyses. M.F., S.H., R.S., and L.C. interpreted the data. M.F. wrote the manuscript with the help of S.H. All authors read and approved the manuscript.

## Declaration of interests

R.S. received speaker’s honorary from AstraZeneca and Roche. All other authors declare no competing interests.

**Supplemental Information** is available for this paper.

---

**Supplementary Figure 1. (A)** UCSC genome browser images displaying publicly available p53 binding signals from Nutlin-3a treated U2OS (42) and HCT116 (43) cells (obtained from CistromeDB (40)) at the *RFX7*, *RFX5*, and *RFX1* gene loci. **(B)** BETA analysis (39) of RFX7 *trans*-activator/repressor function using ChIP-seq and RNA-seq data from U2OS, HCT116, and RPE-1 cells. Based on a correlation between binding proximity to a gene’s TSS and the gene’s differential expression, the BETA analysis tests whether a given TF functions as an activator and/or a repressor of gene expression. Data from two out of three cell lines significantly identified RFX7 to function as a *trans*-activator.

**Supplementary Figure 2.** Heatmap of genes significantly (FDR < 0.01) down-regulated (log_2_FC < -0.25) by siRFX7, up-regulated (log_2_FC > 0.25) by Nutlin-3a (N3A) in siControl treated cells, not bound by RFX7 but with reduced Nutlin-3a-dependent regulation in siRFX7 treated cells. The top 100 genes are displayed ranked by the difference of mean Z-score Nutlin-3a/DMSO difference between siRFX7 and siControl treated cells. *RFX7* is displayed as knockdown control. *CDKN1A* and *MDM2* are direct p53 targets not affected by RFX7.

**Supplementary Figure 3.** Distribution of **(A)** gender, **(B)** age, and **(C)** cancer type in the patient groups. Complement to Figure 6C. **(A)** Statistical significance tested through a pairwise non-parametric Wilcoxon test.

## Reference List

1. Gajiwala,K.S., Chen,H., Cornille,F., Roques,B.P., Reith,W., Mach,B. and Burley,S.K. (2000) Structure of the winged-helix protein hRFX1 reveals a new mode of DNA binding. Nature, 403, 916–921.

2. Sugiaman-Trapman,D., Vitezic,M., Jouhilahti,E.-M., Mathelier,A., Lauter,G., Misra,S., Daub,C.O., Kere,J. and Swoboda,P. (2018) Characterization of the human RFX transcription factor family by regulatory and target gene analysis. BMC Genomics, 19, 181.

3. Aftab,S., Semenec,L., Chu,J.S.-C. and Chen,N. (2008) Identification and characterization of novel human tissue-specific RFX transcription factors. BMC Evol. Biol., 8, 226.

4. López,C., Kleinheinz,K., Aukema,S.M., Rohde,M., Bernhart,S.H., Hübschmann,D., Wagener,R., Toprak,U.H., Raimondi,F., Kreuz,M., et al. (2019) Genomic and transcriptomic changes complement each other in the pathogenesis of sporadic Burkitt lymphoma. Nat. Commun., 10, 1459.

5. Grande,B.M., Gerhard,D.S., Jiang,A., Griner,N.B., Abramson,J.S., Alexander,T.B., Allen,H., Ayers,L.W., Bethony,J.M., Bhatia,K., et al. (2019) Genome-wide discovery of somatic coding and noncoding mutations in pediatric endemic and sporadic Burkitt lymphoma. Blood, 133, 1313.

6. Berndt,S.I., Skibola,C.F., Joseph,V., Camp,N.J., Nieters,A., Wang,Z., Cozen,W., Monnereau,A., Wang,S.S., Kelly,R.S., et al. (2013) Genome-wide association study identifies multiple risk loci for chronic lymphocytic leukemia. Nat. Genet., 45, 868–876.

7. . Speedy,H.E., Di Bernardo,M.C., Sava,G.P., Dyer,M.J.S., Holroyd,A., Wang,Y., Sunter,N.J., Mansouri,L., Juliusson,G., Smedby,K.E., et al. (2014) A genome-wide association study identifies multiple susceptibility loci for chronic lymphocytic leukemia. Nat. Genet., 46, 56–60.

8. Crowther-Swanepoel,D., Broderick,P., Di Bernardo,M.C., Dobbins,S.E., Torres,M., Mansouri,M., Ruiz-Ponte,C., Enjuanes,A., Rosenquist,R., Carracedo,A., et al. (2010) Common variants at 2q37.3, 8q24.21, 15q21.3 and 16q24.1 influence chronic lymphocytic leukemia risk. Nat. Genet., 42, 132–136.

9. . Weber,J., de la Rosa,J., Grove,C.S., Schick,M., Rad,L., Baranov,O., Strong,A., Pfaus,A., Friedrich,M.J., Engleitner,T., et al. (2019) PiggyBac transposon tools for recessive screening identify B-cell lymphoma drivers in mice. Nat. Commun., 10, 1415.

10. Bullinger,L., Krönke,J., Schön,C., Radtke,I., Urlbauer,K., Botzenhardt,U., Gaidzik,V., Carió,A., Senger,C., Schlenk,R.F., et al. (2010) Identification of acquired copy number alterations and uniparental disomies in cytogenetically normal acute myeloid leukemia using high-resolution single-nucleotide polymorphism analysis. Leukemia, 24, 438–449.

11. Rusiniak,M.E., Kunnev,D., Freeland,A., Cady,G.K. and Pruitt,S.C. (2012) Mcm2 deficiency results in short deletions allowing high resolution identification of genes contributing to lymphoblastic lymphoma. Oncogene, 31, 4034–4044.

12. Rogers,L.M., Olivier,A.K., Meyerholz,D.K. and Dupuy,A.J. (2013) Adaptive Immunity Does Not Strongly Suppress Spontaneous Tumors in a Sleeping Beauty Model of Cancer. J. Immunol., 190, 4393–4399.

13. Shungin,D., Winkler,T.W., Croteau-Chonka,D.C., Ferreira,T., Locke,A.E., Mägi,R., Strawbridge,R.J., Pers,T.H., Fischer,K., Justice,A.E., et al. (2015) New genetic loci link adipose and insulin biology to body fat distribution. Nature, 518, 187–196.

14. Kim,D., Basile,A.O., Bang,L., Horgusluoglu,E., Lee,S., Ritchie,M.D., Saykin,A.J. and Nho,K. (2017) Knowledge-driven binning approach for rare variant association analysis: application to neuroimaging biomarkers in Alzheimer’s disease. BMC Med. Inform. Decis. Mak., 17, 61.

15. Harris,H.K., Nakayama,T., Lai,J., Zhao,B., Argyrou,N., Gubbels,C.S., Soucy,A., Genetti,C.A., Suslovitch,V., Rodan,L.H., et al. (2021) Disruption of RFX family transcription factors causes autism, attention-deficit/hyperactivity disorder, intellectual disability, and dysregulated behavior. Genet. Med., 10.1038/s41436-021-01114-z.

16. Manojlovic,Z., Earwood,R., Kato,A., Stefanovic,B. and Kato,Y. (2014) RFX7 is required for the formation of cilia in the neural tube. Mech. Dev., 132, 28–37.

17. Castro,W., Chelbi,S.T., Niogret,C., Ramon-Barros,C., Welten,S.P.M., Osterheld,K., Wang,H., Rota,G., Morgado,L., Vivier,E., et al. (2018) The transcription factor Rfx7 limits metabolism of NK cells and promotes their maintenance and immunity. Nat. Immunol., 19, 809–820.

18. Fischer,M. (2017) Census and evaluation of p53 target genes. Oncogene, 36, 3943–3956.

19. Sammons,M.A., Nguyen,T.-A.T., McDade,S.S. and Fischer,M. (2020) Tumor suppressor p53: from engaging DNA to target gene regulation. Nucleic Acids Res., 48, 8848–8869.

20. Fischer,M., Grossmann,P., Padi,M. and DeCaprio,J.A. (2016) Integration of TP53, DREAM, MMB-FOXM1 and RB-E2F target gene analyses identifies cell cycle gene regulatory networks. Nucleic Acids Res., 44, 6070–6086.

21. Fischer,M., Quaas,M., Steiner,L. and Engeland,K. (2016) The p53-p21-DREAM-CDE/CHR pathway regulates G2/M cell cycle genes. Nucleic Acids Res., 44, 164–174.

22. Uxa,S., Bernhart,S.H., Mages,C.F.S., Fischer,M., Kohler,R., Hoffmann,S., Stadler,P.F., Engeland,K. and Müller,G.A. (2019) DREAM and RB cooperate to induce gene repression and cell-cycle arrest in response to p53 activation. Nucleic Acids Res., 47, 9087–9103.

23. Schade,A.E., Fischer,M. and DeCaprio,J.A. (2019) RB, p130 and p107 differentially repress G1/S and G2/M genes after p53 activation. Nucleic Acids Res., 47, 11197–11208.

24. Baum,N., Schiene-Fischer,C., Frost,M., Schumann,M., Sabapathy,K., Ohlenschläger,O., Grosse,F. and Schlott,B. (2009) The prolyl cis/trans isomerase cyclophilin 18 interacts with the tumor suppressor p53 and modifies its functions in cell cycle regulation and apoptosis. Oncogene, 28, 3915–3925.

25. Bolger,A.M., Lohse,M. and Usadel,B. (2014) Trimmomatic: a flexible trimmer for Illumina sequence data. Bioinformatics, 30, 2114–2120.

26. Martin,M. (2011) Cutadapt removes adapter sequences from high-throughput sequencing reads. EMBnet.journal, 17, 10.

27. Song,L. and Florea,L. (2015) Rcorrector: efficient and accurate error correction for Illumina RNA-seq reads. Gigascience, 4, 48.

28. Kopylova,E., Noé,L. and Touzet,H. (2012) SortMeRNA: Fast and accurate filtering of ribosomal RNAs in metatranscriptomic data. Bioinformatics, 28, 3211–3217.

29. Cunningham,F., Achuthan,P., Akanni,W., Allen,J., Amode,M.R., Armean,I.M., Bennett,R., Bhai,J., Billis,K., Boddu,S., et al. (2019) Ensembl 2019. Nucleic Acids Res., 47, D745–D751.

30. Hoffmann,S., Otto,C., Kurtz,S., Sharma,C.M., Khaitovich,P., Vogel,J., Stadler,P.F. and Hackermüller,J. (2009) Fast Mapping of Short Sequences with Mismatches, Insertions and Deletions Using Index Structures. PLoS Comput. Biol., 5, e1000502.

31. Hoffmann,S., Otto,C., Doose,G., Tanzer,A., Langenberger,D., Christ,S., Kunz,M., Holdt,L.M., Teupser,D., Hackermüller,J., et al. (2014) A multi-split mapping algorithm for circular RNA, splicing, trans-splicing and fusion detection. Genome Biol., 15, R34.

32. Li,H., Handsaker,B., Wysoker,A., Fennell,T., Ruan,J., Homer,N., Marth,G., Abecasis,G. and Durbin,R. (2009) The Sequence Alignment/Map format and SAMtools. Bioinformatics, 25, 2078–2079.

33. Feng,J., Liu,T., Qin,B., Zhang,Y. and Liu,X.S. (2012) Identifying ChIP-seq enrichment using MACS. Nat. Protoc., 7, 1728–40.

34. Quinlan,A.R. and Hall,I.M. (2010) BEDTools: A flexible suite of utilities for comparing genomic features. Bioinformatics, 26, 841–842.

35. Amemiya,H.M., Kundaje,A. and Boyle,A.P. (2019) The ENCODE Blacklist: Identification of Problematic Regions of the Genome. Sci. Rep., 9, 1–5.

36. Heinz,S., Benner,C., Spann,N., Bertolino,E., Lin,Y.C., Laslo,P., Cheng,J.X., Murre,C., Singh,H. and Glass,C.K. (2010) Simple combinations of lineage-determining transcription factors prime cis-regulatory elements required for macrophage and B cell identities. Mol. Cell, 38, 576–89.

37. Siepel,A., Bejerano,G., Pedersen,J.S., Hinrichs,A.S., Hou,M., Rosenbloom,K., Clawson,H., Spieth,J., Hillier,L.W., Richards,S., et al. (2005) Evolutionarily conserved elements in vertebrate, insect, worm, and yeast genomes. Genome Res., 15, 1034–1050.

38. Liu,T., Ortiz,J.A., Taing,L., Meyer,C.A., Lee,B., Zhang,Y., Shin,H., Wong,S.S., Ma,J., Lei,Y., et al. (2011) Cistrome: an integrative platform for transcriptional regulation studies. Genome Biol., 12, R83.

39. Wang,S., Sun,H., Ma,J., Zang,C., Wang,C., Wang,J., Tang,Q., Meyer,C.A., Zhang,Y. and Liu,X.S. (2013) Target analysis by integration of transcriptome and ChIP-seq data with BETA. Nat. Protoc., 8, 2502–2515.

40. Zheng,R., Wan,C., Mei,S., Qin,Q., Wu,Q., Sun,H., Chen,C.-H., Brown,M., Zhang,X., Meyer,C.A., et al. (2019) Cistrome Data Browser: expanded datasets and new tools for gene regulatory analysis. Nucleic Acids Res., 47, D729–D735.

41. Ramírez,F., Ryan,D.P., Grüning,B., Bhardwaj,V., Kilpert,F., Richter,A.S., Heyne,S., Dündar,F. and Manke,T. (2016) deepTools2: a next generation web server for deep-sequencing data analysis. Nucleic Acids Res., 44, W160–W165.

42. Menendez,D., Nguyen,T.A., Freudenberg,J.M., Mathew,V.J., Anderson,C.W., Jothi,R. and Resnick,M.A. (2013) Diverse stresses dramatically alter genome-wide p53 binding and transactivation landscape in human cancer cells. Nucleic Acids Res., 41, 7286–7301.

43. Andrysik,Z., Galbraith,M.D., Guarnieri,A.L., Zaccara,S., Sullivan,K.D., Pandey,A., MacBeth,M., Inga,A. and Espinosa,J.M. (2017) Identification of a core TP53 transcriptional program with highly distributed tumor suppressive activity. Genome Res., 27, 1645–1657.

44. Liao,Y., Smyth,G.K. and Shi,W. (2014) FeatureCounts: An efficient general purpose program for assigning sequence reads to genomic features. Bioinformatics, 30, 923–930.

45. Wang,L., Wang,S. and Li,W. (2012) RSeQC: quality control of RNA-seq experiments. Bioinformatics, 28, 2184–2185.

46. Love,M.I., Huber,W. and Anders,S. (2014) Moderated estimation of fold change and dispersion for RNA-seq data with DESeq2. Genome Biol., 15, 550.

47. Ramirez,R.N., El-Ali,N.C., Mager,M.A., Wyman,D., Conesa,A. and Mortazavi,A. (2017) Dynamic Gene Regulatory Networks of Human Myeloid Differentiation. Cell Syst., 4, 416–429.e3.

48. Schira-Heinen,J., Czapla,A., Hendricks,M., Kloetgen,A., Wruck,W., Adjaye,J., Kögler,G., Werner Müller,H., Stühler,K. and Trompeter,H.-I. (2020) Functional omics analyses reveal only minor effects of microRNAs on human somatic stem cell differentiation. Sci. Rep., 10, 3284.

49. Jacob,A., Morley,M., Hawkins,F., McCauley,K.B., Jean,J.C., Heins,H., Na,C.-L., Weaver,T.E., Vedaie,M., Hurley,K., et al. (2017) Differentiation of Human Pluripotent Stem Cells into Functional Lung Alveolar Epithelial Cells. Cell Stem Cell, 21, 472–488.e10.

50. . Janky,R., Verfaillie,A., Imrichová,H., van de Sande,B., Standaert,L., Christiaens,V., Hulselmans,G., Herten,K., Naval Sanchez,M., Potier,D., et al. (2014) iRegulon: From a Gene List to a Gene Regulatory Network Using Large Motif and Track Collections. PLoS Comput. Biol., 10, e1003731.

51. Tsherniak,A., Vazquez,F., Montgomery,P.G., Weir,B.A., Kryukov,G., Cowley,G.S., Gill,S., Harrington,W.F., Pantel,S., Krill-Burger,J.M., et al. (2017) Defining a Cancer Dependency Map. Cell, 170, 564–576.e16.

52. McFarland,J.M., Ho,Z. V., Kugener,G., Dempster,J.M., Montgomery,P.G., Bryan,J.G., Krill-Burger,J.M., Green,T.M., Vazquez,F., Boehm,J.S., et al. (2018) Improved estimation of cancer dependencies from large-scale RNAi screens using model-based normalization and data integration. Nat. Commun., 9, 4610.

53. Colaprico,A., Silva,T.C., Olsen,C., Garofano,L., Cava,C., Garolini,D., Sabedot,T.S., Malta,T.M., Pagnotta,S.M., Castiglioni,I., et al. (2016) TCGAbiolinks: An R/Bioconductor package for integrative analysis of TCGA data. Nucleic Acids Res., 44, e71.

54. Barbie,D.A., Tamayo,P., Boehm,J.S., Kim,S.Y., Moody,S.E., Dunn,I.F., Schinzel,A.C., Sandy,P., Meylan,E., Scholl,C., et al. (2009) Systematic RNA interference reveals that oncogenic KRAS-driven cancers require TBK1. Nature, 462, 108–112.

55. Barrett,T., Wilhite,S.E., Ledoux,P., Evangelista,C., Kim,I.F., Tomashevsky,M., Marshall,K.A., Phillippy,K.H., Sherman,P.M., Holko,M., et al. (2013) NCBI GEO: Archive for functional genomics data sets - Update. Nucleic Acids Res., 41, D991–5.

56. ENCODE Project Consortium (2012) An integrated encyclopedia of DNA elements in the human genome. Nature, 489, 57–74.

57. Hilliard,A., Hilliard,B., Zheng,S.-J., Sun,H., Miwa,T., Song,W., Göke,R. and Chen,Y.H. (2006) Translational Regulation of Autoimmune Inflammation and Lymphoma Genesis by Programmed Cell Death 4. J. Immunol., 177, 8095–8102.

58. He,X., Zhu,Z., Johnson,C., Stoops,J., Eaker,A.E., Bowen,W. and DeFrances,M.C. (2008) PIK3IP1, a negative regulator of PI3K, suppresses the development of hepatocellular carcinoma. Cancer Res., 68, 5591–8.

59. Gaviraghi,M., Vivori,C., Pareja Sanchez,Y., Invernizzi,F., Cattaneo,A., Santoliquido,B.M., Frenquelli,M., Segalla,S., Bachi,A., Doglioni,C., et al. (2018) Tumor suppressor PNRC1 blocks rRNA maturation by recruiting the decapping complex to the nucleolus. EMBO J., 37, e99179.

60. Schaub,F.X., Dhankani,V., Berger,A.C., Trivedi,M., Richardson,A.B., Shaw,R., Zhao,W., Zhang,X., Ventura,A., Liu,Y., et al. (2018) Pan-cancer Alterations of the MYC Oncogene and Its Proximal Network across the Cancer Genome Atlas. Cell Syst., 6, 282–300.e2.

61. Vassilev,L.T., Vu,B.T., Graves,B., Carvajal,D., Podlaski,F., Filipovic,Z., Kong,N., Kammlott,U., Lukacs,C., Klein,C., et al. (2004) In vivo activation of the p53 pathway by small-molecule antagonists of MDM2. Science, 303, 844–8.

62. Fischer,B.A., Chelbi,S.T. and Guarda,G. (2020) Regulatory Factor X 7 and its Potential Link to Lymphoid Cancers. Trends in Cancer, 6, 6–9.

63. Nardini,M., Gnesutta,N., Donati,G., Gatta,R., Forni,C., Fossati,A., Vonrhein,C., Moras,D., Romier,C., Bolognesi,M., et al. (2013) Sequence-specific transcription factor NF-Y displays histone-like DNA binding and H2B-like ubiquitination. Cell, 152, 132–143.

64. Reith,W., LeibundGut-Landmann,S. and Waldburger,J.-M. (2005) Regulation of MHC class II gene expression by the class II transactivator. Nat. Rev. Immunol., 5, 793–806.

65. Chen,X., Cao,Q., Liao,R., Wu,X., Xun,S., Huang,J. and Dong,C. (2019) Loss of ABAT-Mediated GABAergic System Promotes Basal-Like Breast Cancer Progression by Activating Ca2+-NFAT1 Axis. Theranostics, 9, 34–47.

66. Bernaudo,S., Salem,M., Qi,X., Zhou,W., Zhang,C., Yang,W., Rosman,D., Deng,Z., Ye,G., Yang,B.B., et al. (2016) Cyclin G2 inhibits epithelial-to-mesenchymal transition by disrupting Wnt/β-catenin signaling. Oncogene, 35, 4816–27.

67. Nagata,E., Luo,H.R., Saiardi,A., Bae,B. II, Suzuki,N. and Snyder,S.H. (2005) Inositol hexakisphosphate kinase-2, a physiologic mediator of cell death. J. Biol. Chem., 280, 1634–1640.

68. Li,F., Sun,Q., Liu,K., Zhang,L., Lin,N., You,K., Liu,M., Kon,N., Tian,F., Mao,Z., et al. (2020) OTUD5 cooperates with TRIM25 in transcriptional regulation and tumor progression via deubiquitination activity. Nat. Commun., 11, 4184.

69. Wittschieben,J.P., Patil,V., Glushets,V., Robinson,L.J., Kusewitt,D.F. and Wood,R.D. (2010) Loss of DNA polymerase zeta enhances spontaneous tumorigenesis. Cancer Res., 70, 2770–8.

70. Gawrzak,S., Rinaldi,L., Gregorio,S., Arenas,E.J., Salvador,F., Urosevic,J., Figueras-Puig,C., Rojo,F., del Barco Barrantes,I., Cejalvo,J.M., et al. (2018) MSK1 regulates luminal cell differentiation and metastatic dormancy in ER+ breast cancer. Nat. Cell Biol., 20, 211–221.

71. Yoshida,Y., Nakamura,T., Komoda,M., Satoh,H., Suzuki,T., Tsuzuku,J.K., Miyasaka,T., Yoshida,E.H., Umemori,H., Kunisaki,R.K., et al. (2003) Mice lacking a transcriptional corepressor Tob are predisposed to cancer. Genes Dev., 17, 1201–1206.

72. Yu,J., Ershler,M., Yu,L., Wei,M., Hackanson,B., Yokohama,A., Mitsui,T., Liu,C.C.G., Mao,H., Liu,S., et al. (2009) TSC-22 contributes to hematopoietic precursor cell proliferation and repopulation and is epigenetically silenced in large granular lymphocyte leukemia. Blood, 113, 5558–5567.

73. Epping,M.T., Lunardi,A., Nachmani,D., Castillo-Martin,M., Thin,T.H., Cordon-Cardo,C. and Pandolfi,P.P. (2015) TSPYL2 is an essential component of the REST/NRSF transcriptional complex for TGFβ signaling activation. Cell Death Differ., 22, 1353–1362.

74. Hoadley,K.A., Yau,C., Hinoue,T., Wolf,D.M., Lazar,A.J., Drill,E., Shen,R., Taylor,A.M., Cherniack,A.D., Thorsson,V., et al. (2018) Cell-of-Origin Patterns Dominate the Molecular Classification of 10,000 Tumors from 33 Types of Cancer. Cell, 173, 291–304.e6.

75. Whitfield,M.L., George,L.K., Grant,G.D. and Perou,C.M. (2006) Common markers of proliferation. Nat. Rev. Cancer, 6, 99–106.

76. Hanahan,D. and Weinberg,R.A. (2000) The Hallmarks of Cancer. Cell, 100, 57–70.

77. Hurlin,P.J., Quéva,C., Koskinen,P.J., Steingrímsson,E., Ayer,D.E., Copeland,N.G., Jenkins,N.A. and Eisenman,R.N. (1995) Mad3 and Mad4: novel Max-interacting transcriptional repressors that suppress c-myc dependent transformation and are expressed during neural and epidermal differentiation. EMBO J., 14, 5646–59.

78. Yeo,C.Q.X., Alexander,I., Lin,Z., Lim,S., Aning,O.A., Kumar,R., Sangthongpitag,K., Pendharkar,V., Ho,V.H.B. and Cheok,C.F. (2016) P53 Maintains Genomic Stability by Preventing Interference between Transcription and Replication. Cell Rep., 15, 132–146.

79. Ried,J.S., Jeff M.J., Chu,A.Y., Bragg-Gresham,J.L., van Dongen,J., Huffman,J.E., Ahluwalia,T.S., Cadby,G., Eklund,N., Eriksson,J., et al. (2016) A principal component meta-analysis on multiple anthropometric traits identifies novel loci for body shape. Nat. Commun., 7, 13357.

