## Supplementary figures and images for "Transcription factor RFX7 governs a tumor suppressor network in response to p53 and stress"

### Supplementary Figure 1

Coronel *et al.*, Supplementary Figure 1

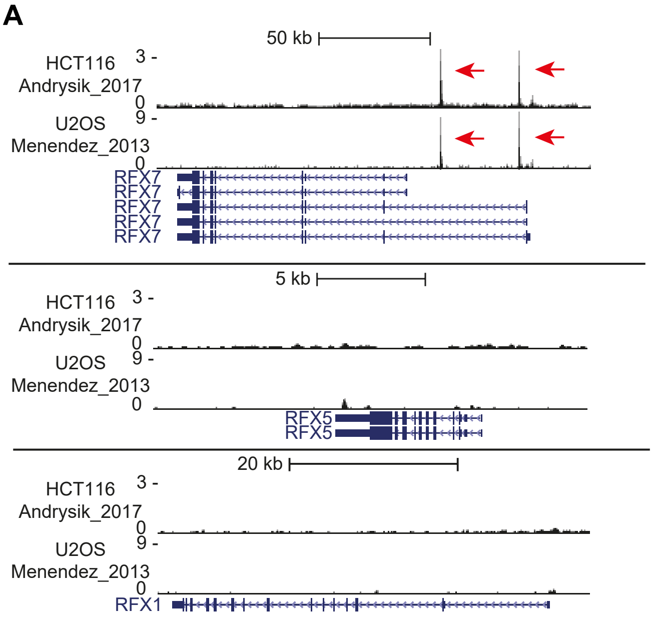


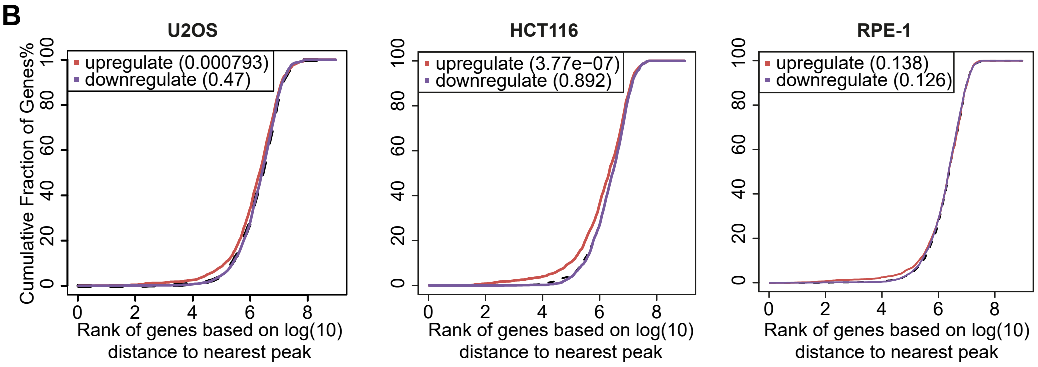

### Supplementary Figure 2

Coronel *et al.*, Supplementary Figure 2

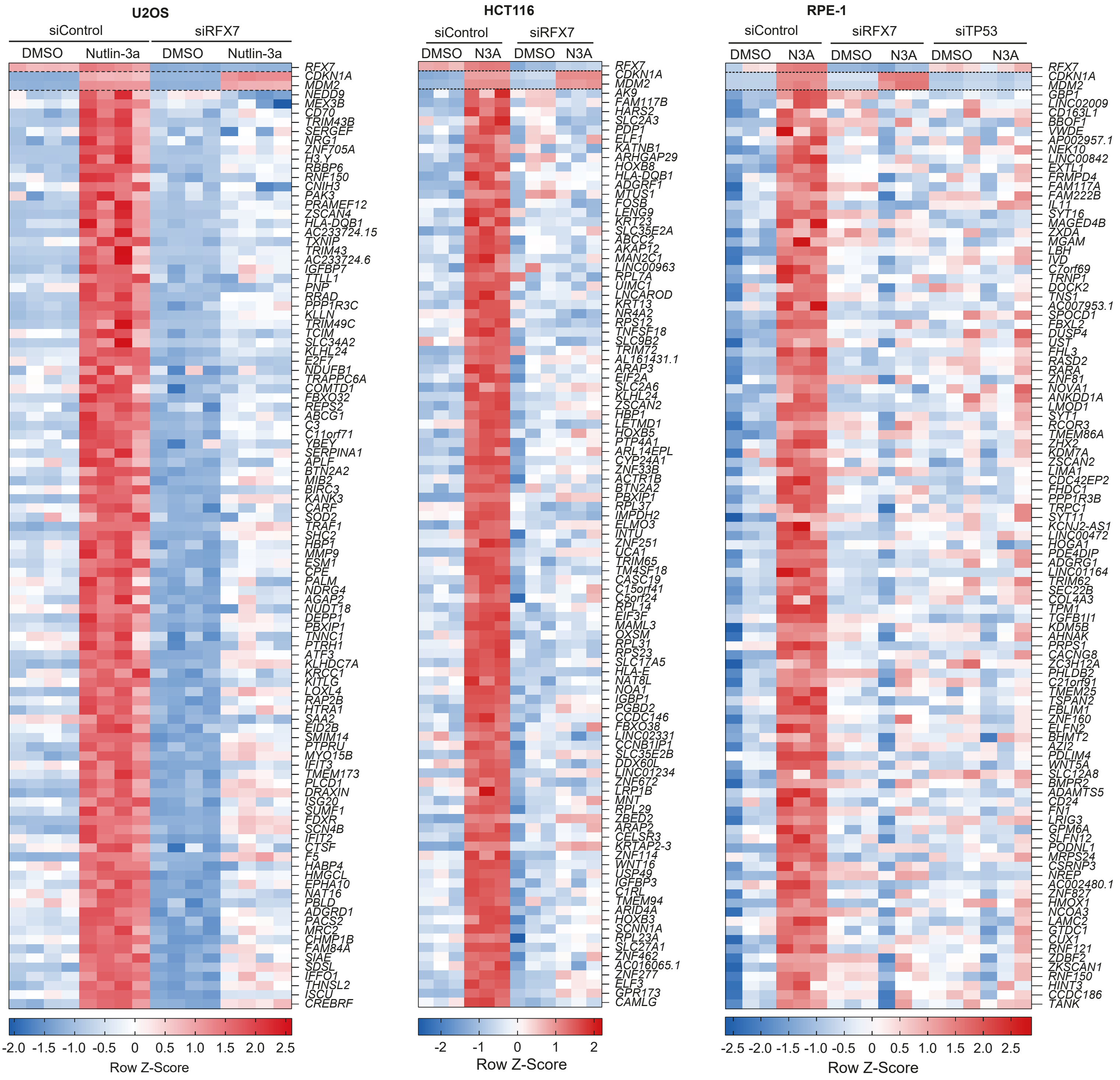

### Supplementary Figure 3

Coronel *et al.*, Supplementary Figure 3

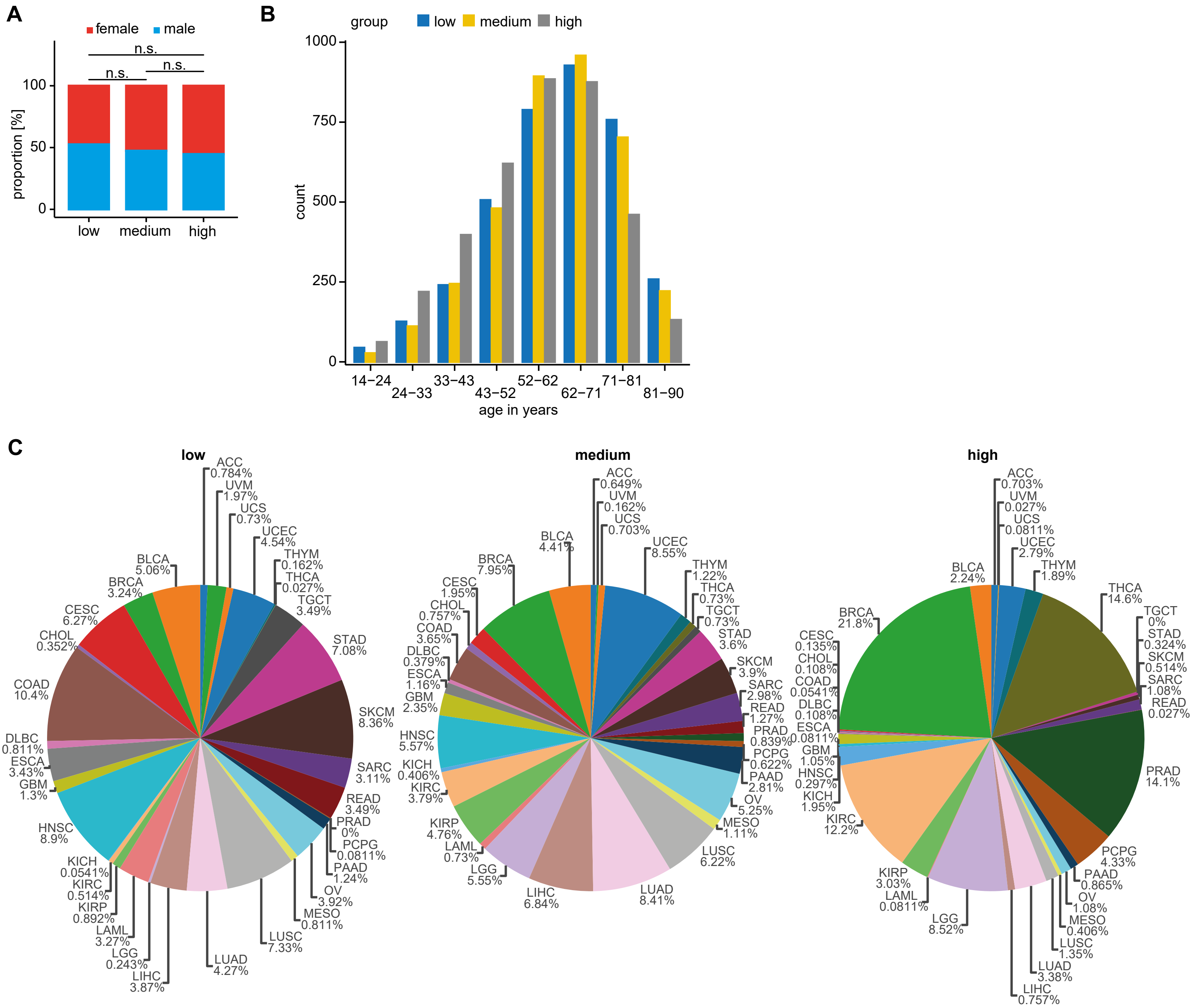
